# Recurrent Connections in the Primate Ventral Visual Stream Mediate a Tradeoff Between Task Performance and Network Size During Core Object Recognition

**DOI:** 10.1101/2021.02.17.431717

**Authors:** Aran Nayebi, Javier Sagastuy-Brena, Daniel M. Bear, Kohitij Kar, Jonas Kubilius, Surya Ganguli, David Sussillo, James J. DiCarlo, Daniel L. K. Yamins

## Abstract

The computational role of the abundant feedback connections in the ventral visual stream (VVS) is unclear, enabling humans and non-human primates to effortlessly recognize objects across a multitude of viewing conditions. Prior studies have augmented feedforward convolutional neural networks (CNNs) with recurrent connections to study their role in visual processing; however, often these recurrent networks are optimized directly on neural data or the comparative metrics used are undefined for standard feedforward networks that lack these connections. In this work, we develop *task-optimized* convolutional recurrent (ConvRNN) network models that more correctly mimic the timing and gross neuroanatomy of the ventral pathway. Properly chosen intermediate-depth ConvRNN circuit architectures, which incorporate mechanisms of feedforward bypassing and recurrent gating, can achieve high performance on a core recognition task, comparable to that of much deeper feedforward networks. We then develop methods that allow us to compare both CNNs and ConvRNNs to fine-grained measurements of primate categorization behavior and neural response trajectories across thousands of stimuli. We find that high performing ConvRNNs provide a better match to this data than feedforward networks of any depth, predicting the precise timings at which each stimulus is behaviorally decoded from neural activation patterns. Moreover, these ConvRNN circuits consistently produce quantitatively accurate predictions of neural dynamics from V4 and IT across the entire stimulus presentation. In fact, we find that the highest performing ConvRNNs, which best match neural and behavioral data, also achieve a strong Pareto-tradeoff between task performance and overall network size. Taken together, our results suggest the functional purpose of recurrence in the ventral pathway is to fit a high performing network in cortex, attaining computational power through temporal rather than spatial complexity.

## 1 Introduction

The visual system of the brain must discover meaningful patterns in a complex physical world^1^. Within 200*ms*, primates can quickly identify objects despite changes in position, pose, contrast, background, foreground, and many other factors from one occasion to the next: a behavior known as “core object recognition”^2,3^. It is known that the ventral visual stream (VVS) underlies this ability by transforming the retinal image of an object into a new internal representation, in which high-level properties, such as object identity and category, are more explicit^3^.

Non-trivial dynamics result from a ubiquity of recurrent connections in the VVS, including synapses that facilitate or depress, dense local recurrent connections within each cortical region, and long-range connections between different regions, such as feedback from higher to lower visual cortex^4^. Furthermore, the behavioral roles of recurrence and dynamics in the visual system are not well understood. Several conjectures are that recurrence “fills in” missing data,^5,6,7,8^ such as object parts occluded by other objects; that it “sharpens” representations by top-down attentional feature refinement, allowing for easier decoding of certain stimulus properties or performance of certain tasks^4,9,10,11,12^; that it allows the brain to “predict” future stimuli (such as the frames of a movie)^13,14,15^; or that recurrence “extends” a feedforward computation, reflecting the fact that an unrolled recurrent network is equivalent to a deeper feedforward network that conserves on neurons (and learnable parameters) by repeating transformations several times^16,17,18,7,19,20^. Formal computational models are needed to test these hypotheses: if optimizing a model for a certain task leads to accurate predictions of neural dynamics, then that task may be a primary reason those dynamics occur in the brain.

We therefore broaden the method of goal-driven modeling from solving tasks with feedforward CNNs^21^ or RNNs^22^ to explain dynamics in the primate visual system, building convolutional recurrent neural networks (ConvRNNs), depicted in Figure 1. There has been substantial prior work in this domain^16,10,17,19,23,20^, which we go beyond in several important ways.

**Figure 1:**
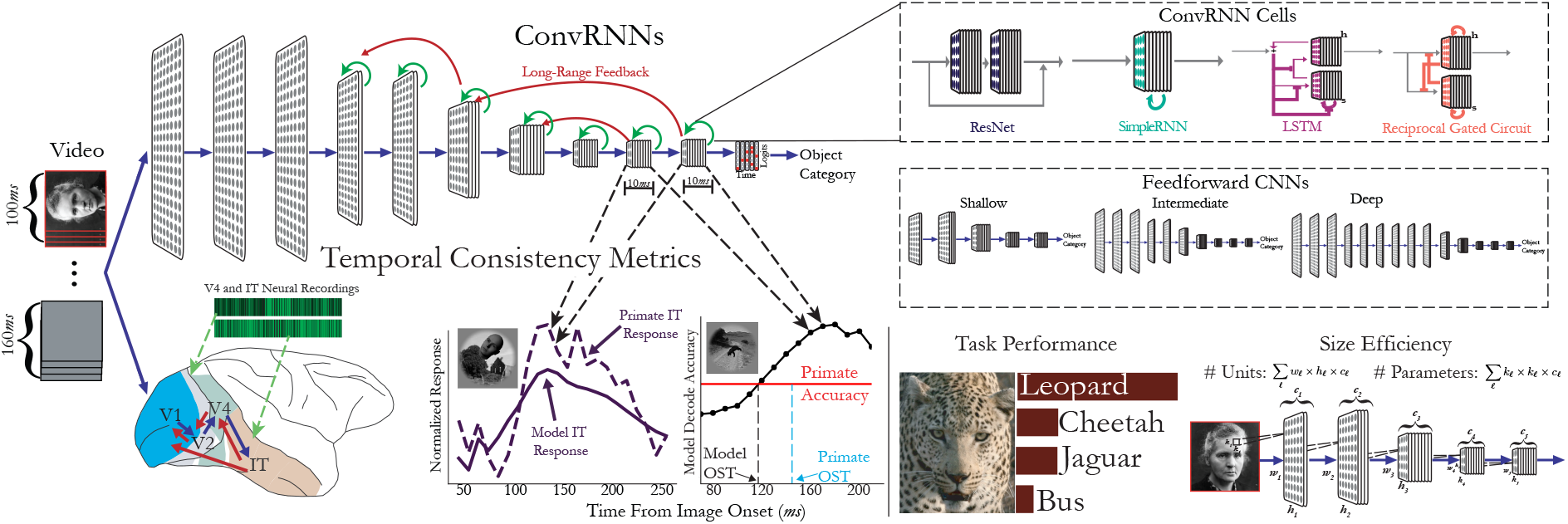
ConvRNNs as models of the primate ventral visual stream. *Performance-optimized recurrence*. Convolutional recurrent networks (ConvRNNs) have a combination of local recurrent circuits (green) and long-range feedback connections (red) added on top of a feedforward CNN “BaseNet” backbone (blue). Feedforward CNNs are therefore a special case of ConvRNNs, and we consider a variety of CNNs of varying depths, trained on the ImageNet categorization task. We also perform large-scale evolutionary searches over the local and long-range feedback connections. In addition, we consider particular choices of “light-weight” (in terms of parameter count) decoding strategy that determines the final object category of that image. In our implementation displayed on the top, propagation along each arrow takes one time step (10*ms*) to mimic conduction delays between cortical layers. *Measurements*. From each network class, we measure categorization performance and its size in terms of its parameter and neuron count. *Comparison to neural and behavioral data*. Each stimulus was presented for 100*ms*, followed by a mean gray stimulus interleaved between images, lasting a total of 260*ms*. All images were presented to the models for 10 time steps (corresponding to 100*ms*), followed by a mean gray stimulus for the remaining 15 time steps, to match the image presentation to the primates. We stipulated that units from each multi-unit array must be fit by features from a single model layer, detailed in Section A.6.2. Model features produce a temporally-varying output that can be compared to primate neural dynamics in V4 and IT, as well as temporally-varying behaviors in the form of object solution times (OST).

We show that with a novel choice of layer-local recurrent circuit and long-range feedback connectivity pattern, ConvRNNs can match the performance of much deeper feedforward CNNs on ImageNet but with far fewer units and parameters, as well as a more anatomically consistent number of layers, by extending these computations through time. In fact, such ConvRNNs most accurately explain neural dynamics from V4 and IT across the entirety of stimulus presentation with a temporally-fixed linear mapping, compared to alternative recurrent circuits. Furthermore, we find these suitably-chosen ConvRNN circuit architectures provide a better match to primate behavior in the form of object solution times, compared to feedforward CNNs. We observe that ConvRNNs that attain high task performance but have small overall network size, as measured by number of units, are most consistent with this data – while even the highest-performing but biologically-implausible deep feedforward models are overall a *less consistent* match. In fact, we find a strong Pareto-tradeoff between network size and performance, with ConvRNNs of biologically-plausible intermediate-depth occupying the sweet spot with high performance and a (comparatively) small overall network size. Because we do not fit neural networks end-to-end to neural data (c.f.^23^), but instead show that these outcomes emerge naturally from task performance, our approach enables a normative interpretation of the structural and functional design principles of the model.

Our work is also the first to develop large-scale task-optimized ConvRNNs with biologically-plausible temporal unrolling. Unlike most studies of combinations of convolutional and recurrent networks, which posit a recurrent subnetwork concatenated onto the end of a convolutional backbone^10^, we model local recurrence implanted within ConvRNN layers, and allow long-range feedback between layers. Moreover, we treat each connection in the network – whether feedforward or feedback – as a real temporal object with a biophysical conduction delay (set at ~10*ms*), rather than the typical procedure (e.g. as in^10,17,19^) in which the feedforward component of the network (no matter now deep) operates in one timestep. As a result, our networks can be directly compared with neural and behavioral trajectories at a fine-grained scale limited only by the conduction delay itself.

This level of realism is especially important for establishing what we have found appears to be the main real quantitative advantage of ConvRNNs as biological models, as compared to very deep feedforward networks. In particular, we can define an improved metric for assessing the correctness of the match between a ConvRNN network – thought of as a dynamical system – and the actual trajectories of real neurons. By treating such feedforward networks as ConvRNNs with recurrent connections set to 0, we can map these networks to temporal trajectories as well. As a result, we can directly ask, how much of the neural-behavioral trajectory of real brain data is explicable by very deep feedforward networks? This is an important question because implausibly deep networks have been shown in the literature not only to achieve the highest categorization performance^24^ but also achieve competitive results on matching static (temporally-averaged) neural responses^25^. Due to non-biological temporal unrolling, previous work with comparing such networks to temporal trajectories in neural data^19^ has been forced to unfairly score feedforward networks as total failures, with temporal match score artificially set at 0. With our improved realism, we find (see results section below) that deep feedforward networks actually make quite nontrivial temporal predictions that do explain *some* of the reliable temporal variability of real neurons. In this context, our finding that plausibly-deep ConvRNNs in turn meaningfully outperform these deep feedforward networks on this more fair metric is a strong and nontrivial signal of the actually-better biological match of ConvRNNs as compared to deep feedforward networks.

## 2 Results

### 2.1 An evolutionary architecture search yields specific layer-local recurrent circuits and long-range feedback connectivity patterns that improve task performance and maintain small network size

We first tested whether augmenting CNNs with standard RNN circuits from the machine learning community, SimpleRNNs and LSTMs, could improve performance on ImageNet object recognition (Figure 2a). We found that these recurrent circuits added a small amount of accuracy when introduced into the convolutional layers of a shallow, 6-layer feedforward backbone (“FF” in Figure S1) based off of the AlexNet^26^ architecture, which we will refer to as a “BaseNet” (see Section A.3 for architecture details). However, there were two problems with these resultant recurrent architectures: first, these ConvRNNs did not perform substantially better than parameter-matched, minimally unrolled controls – defined as the minimum number of timesteps after the initial feedforward pass whereby all recurrence connections were engaged at least once. This control comparison suggested that the observed performance gain was due to an increase in the number of unique parameters added by the implanted ConvRNN circuits rather than temporally-extended recurrent computation. Second, making the feedforward model wider or deeper yielded an even larger performance gain than adding these standard RNN circuits, but with fewer parameters. This suggested that standard RNN circuits, although well-suited for a range of temporal tasks, are less well-suited for inclusion within deep CNNs to solve challenging object recognition tasks.

**Figure 2:**
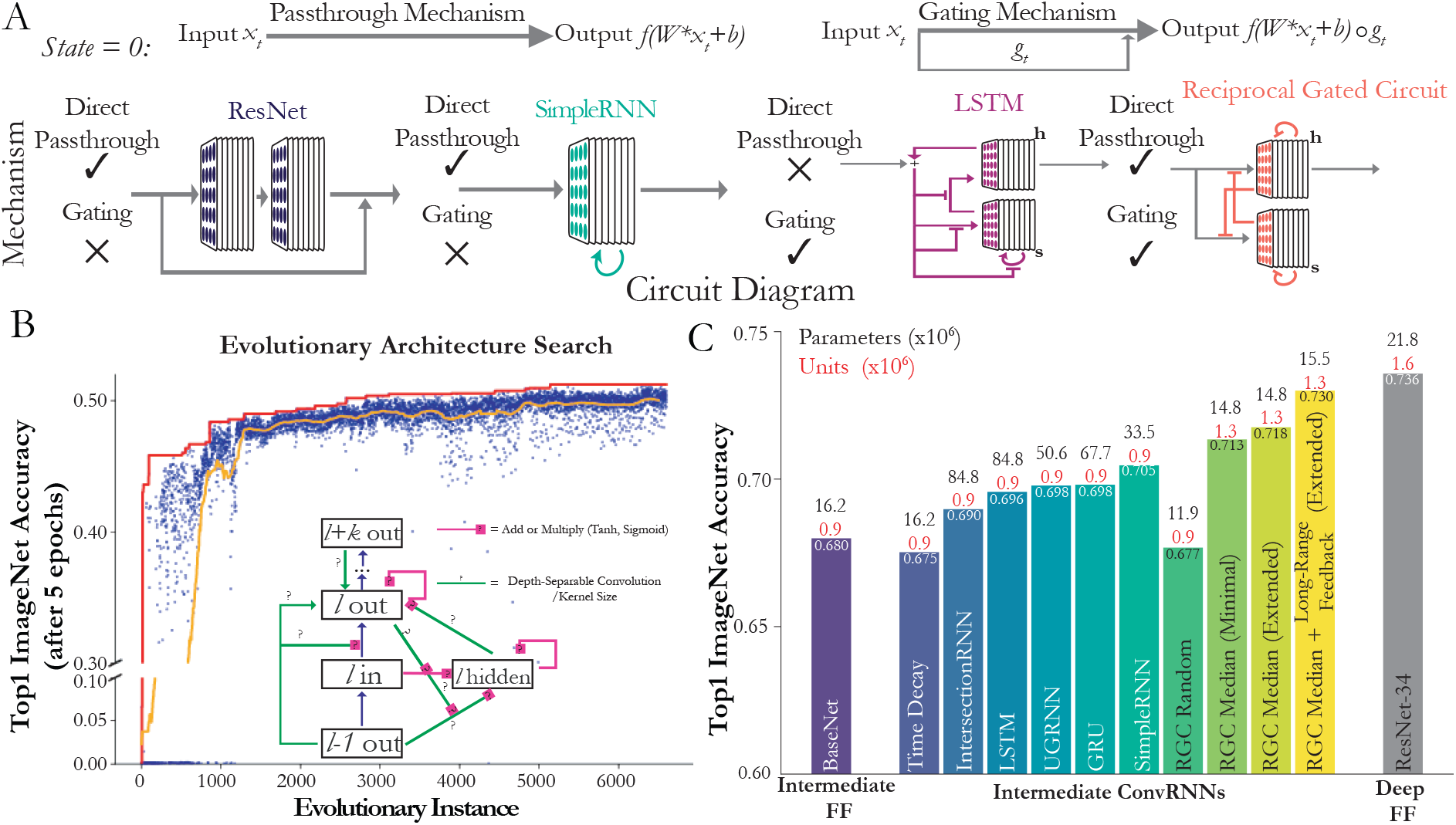
Suitably-chosen intermediate ConvRNN circuits can match the object recognition performance of much deeper feedforward models. **(a) Architectural differences between ConvRNN circuits.** Standard ResNet blocks and SimpleRNN circuits have direct passthrough but not gating. Namely, on the first timestep, the output of a given ConvRNN layer is directly a single linear-nonlinear function of its input, equivalent to that of a feedforward CNN layer (namely, *f* (*W* * *x_t_* + *b*), where *f* is a nonlinear function such as ELU/ReLU and *x_t_* is the input). The LSTM circuit has gating, denoted by T-junctions, but not direct passthrough. The Reciprocal Gated Circuit has both. **(b) ConvRNN circuit search.** Each blue dot represents a model, sampled from hyperparameter space, trained for five epochs. The orange line is the average performance of the last 50 models up to that time. The red line denotes the top performing model at that point in the search. *Search space schematic*: Question marks denote optional connections, which may be conventional or depth-separable convolutions with a choice of kernel size. **(c) Performance of models fully trained on ImageNet.** We compared the performance of an 11-layer feedforward base model (“BaseNet”) modeled after ResNet-18, a control ConvRNN model with trainable time constants (“Time Decay”), along with various other common RNN architectures implanted into this BaseNet, as well as the median Reciprocal Gated Circuit (RGC) model from the search (“RGC Median”) with or without global feedback connectivity, and its minimally-unrolled control (see the first table in Section A.3 for the exact timestep values). The “RGC Random” model was selected randomly from the initial, random phase of the model search. Parameter and unit counts (total number of neurons in the output of each layer) in millions are shown on top of each bar.

We speculated that this was because standard circuits lack a combination of two key properties, each of which on their own have been successful either purely for RNNs or for feedforward CNNs: (1) **Direct passthrough**, where at the first timestep, a zero-initialized hidden state allows feedforward input to pass on to the next layer as a single linear-nonlinear layer just as in a standard feedforward CNN layer (Figure 2a; top left); and (2) **Gating**, in which the value of a hidden state determines how much of the bottom-up input is passed through, retained, or discarded at the next time step (Figure 2a; top right). For example, LSTMs employ gating, but not direct passthrough, as their inputs must pass through several nonlinearities to reach their output; whereas SimpleRNNs do passthrough a zero-initialized hidden state, but do not gate their input (Figure 2a; see Section A.3 for cell equations). Additionally, each of these computations have direct analogies to biological mechanisms – “direct passthrough” would correspond to feedforward processing in time, and “gating” would correspond to adaptation to stimulus statistics across time^27,28^.

We thus implemented recurrent circuits with both features to determine whether they function better than standard circuits within CNNs. One example of such a structure is the “Reciprocal Gated Circuit” (RGC)^29^, which passes through its zero-initialized hidden state and incorporates gating (Figure 2a, bottom right; see Section A.3.7 for the circuit equations). Adding this circuit to the 6-layer BaseNet (“FF”) increased accuracy from 0.51 and 0.53 (“RGC Minimal”, the minimally unrolled, parameter-matched control version) to 0.6 (“RGC Extended”). Moreover, the RGC used substantially fewer parameters than the standard circuits to achieve greater accuracy (Figure S1).

However, it has been shown that different RNN structures can succeed or fail to perform a given task because of differences in trainability rather than differences in capacity^30^. Therefore, we designed an evolutionary search to jointly optimize over both discrete choices of recurrent connectivity patterns as well as continuous choices of learning hyperparameters and weight initializations (search details in Section A.4). While a large-scale search over thousands of convolutional LSTM architectures did yield a better purely gated LSTM-based ConvRNN (“LSTM Opt”), it did not eclipse the performance of the smaller RGC ConvRNN. In fact, applying the same hyperparameter optimization procedure to the RGC ConvRNNs equally increased that architecture class’s performance and further reduced its parameter count (Figure S1, “RGC Opt”).

Therefore, given the promising results with shallower networks, we turned to embedding recurrent circuit motifs into intermediate-depth feedforward networks at scale, whose number of feedforward layers corresponds to the timing of the ventral stream^3^. We then performed an evolutionary search over these resultant intermediate-depth recurrent architectures (Figure 2b). If the primate visual system uses recurrence in lieu of greater network depth to perform object recognition, then a shallower recurrent model with a suitable form of recurrence should achieve recognition accuracy equal to a deeper feedforward model, albeit with temporally-fixed parameters^16^. We therefore tested whether our search had identified such well-adapted recurrent architectures by fully training a representative ConvRNN, the model with the median (across 7000 sampled models) validation accuracy after five epochs of ImageNet training. This median model (“RGC Median”) reached a final ImageNet top-1 validation accuracy nearly equal to a ResNet-34 model with nearly twice as many layers, even though the ConvRNN used only ~ 75% as many parameters. The fully unrolled model from the random phase of the search (“RGC Random”) did not perform substantially better than the BaseNet, though the minimally unrolled control did (Figure 2c). We suspect the improvement of the base intermediate feedforward model over using shallow networks (as in Figure S1) diminishes the difference between the minimal and extended versions of the RGC compared to suitably chosen long-range feedback connections. However, compared to alternative choices of ConvRNN circuits, even the minimally extended RGC significantly outperforms them with fewer parameters and units, indicating the importance of this circuit motif for task performance. This observation suggests that our evolutionary search strategy yielded effective recurrent architectures beyond the initial random phase of the search.

We also considered a control model (“Time Decay”) that produces temporal dynamics by learning time constants on the activations independently at each layer, rather than by learning connectivity between units. In this ConvRNN, unit activations have exponential rather than immediate falloff once feedforward drive ceases. These dynamics could arise, for instance, from single-neuron biophysics (e.g. synaptic depression) rather than interneuronal connections. However, this model did not perform any better than the feedforward BaseNet, implying that ConvRNN performance is not a trivial result of outputting a dynamic time course of responses. We further implanted other more sophisticated forms of ConvRNN circuits into the BaseNet, and while this improved performance over the Time Decay model, it did not outperform the RGC Median ConvRNN despite having many more parameters (Figure 2c). Together, these results demonstrate that the RGC Median ConvRNN uses recurrent computations to subserve object recognition behavior and that particular motifs in its recurrent architecture (Figure S2), found through an evolutionary search, are required for its improved accuracy. Thus, given suitable local recurrent circuits and patterns of long-range feedback connectivity, a physically more compact, temporally-extended ConvRNN can do the same challenging object recognition task as a deeper feedforward CNN.

### 2.2 ConvRNNs better match temporal dynamics of primate behavior than feedforward models

To address whether recurrent processing is engaged during core object recognition behavior, we turn to behavioral data collected from behaving primates. There is a growing body of evidence that current feedforward models fail to accurately capture primate behavior^31,12^. We therefore reasoned that if recurrence is critical to core object recognition behavior, then recurrent networks should be more consistent with suitable measures of primate behavior compared to the feedforward model family. Since the identity of different objects is decoded from the IT population at different times, we considered the first time at which the IT neural decoding accuracy reaches the (pooled) primate behavioral accuracy of a given image, known as the “object solution time (OST)”^12^. Given that our ConvRNNs also have an output at each 10*ms* timebin, the procedure for computing the OST for these models is computed from its “IT-preferred” layers, and we report the “OST consistency”, which we define as the Spearman correlation between the model OSTs and the IT population’s OSTs on the common set of images solved by the given model and IT under the *same* stimulus presentation (see Sections A.6.1 and A.8 for more details).

Unlike our ConvRNNs, which exhibit more biologically plausible temporal dynamics, evaluating the temporal dynamics in feedforward models poses an immediate problem. Given that recurrent networks repeatedly apply nonlinear transformations across time, we can analogously map the layers of a feedforward network to timepoints, observing that a network with *k* distinct layers can produce *k* distinct OSTs in this manner. Thus, the most direct definition of a feedforward model’s OST is to uniformly distribute the timebins between 70-260*ms* across its *k* layers. For very deep feedforward networks such as ResNet-101 and ResNet-152, this number of distinct layers will be as fine-grained as the 10*ms* timebins of the IT responses; however, for most other shallower feedforward networks this will be much coarser. Therefore to enable these feedforward models to be maximally temporally expressive, we *additionally* randomly sample units from consecutive feedforward layers to produce a more graded temporal mapping, depicted in Figure 3a. This graded mapping is ultimately what we use for the feedforward models in Figure 3c, providing the highest OST consistency for that model class^*a*^. Note that for ConvRNNs and very deep feedforward models (ResNet-101 and ResNet-152) whose number of “IT-preferred” layers matches the number of timebins, then the uniform and graded mappings are equivalent, whereby the earliest (in the feedforward hierarchy) layer is matched to the earliest 10*ms* timebin of 70*ms*, and so forth.

**Figure 3:**
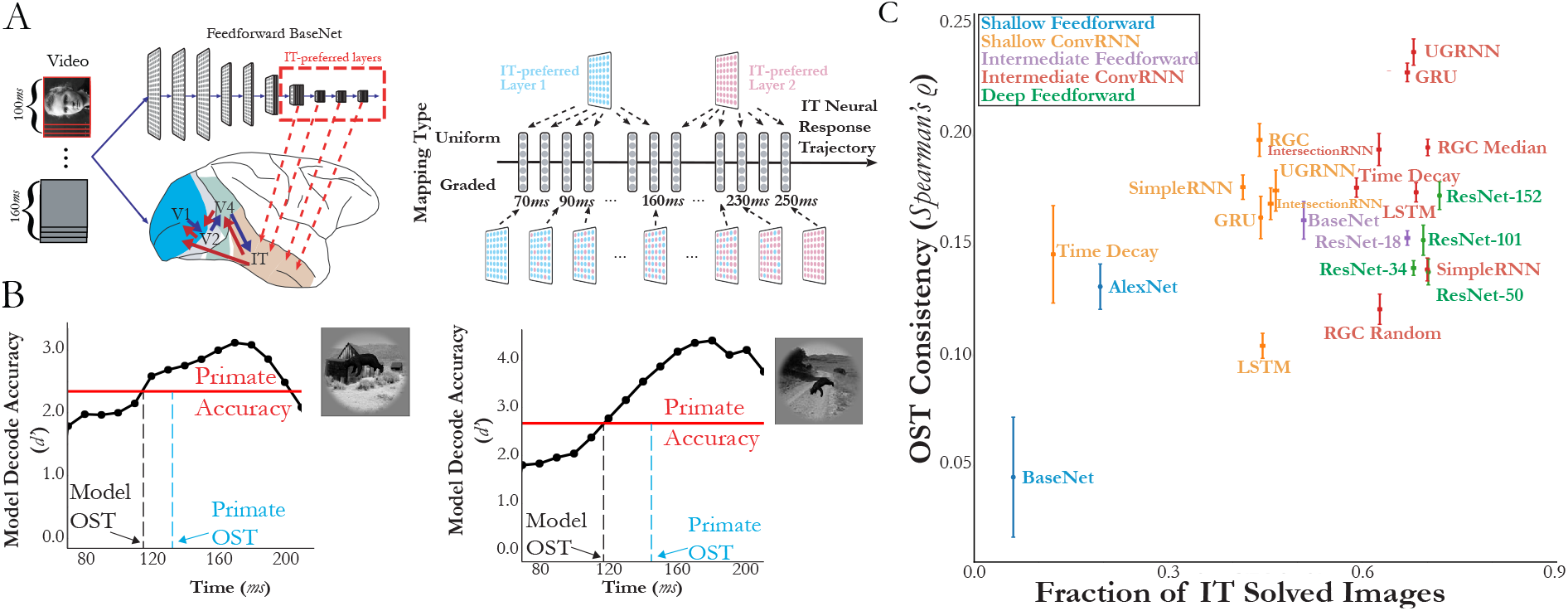
Intermediate ConvRNNs best explain the object solution times (OST) of IT across images. **(a) Comparing to primate OSTs.** *Mapping model layers to timepoints*. In order to compare to primate IT object solution times, namely the first time at which the neural decode accuracy for each image reached the level of the (pooled) primate behavioral accuracy, we first need to define object solution times for models. This procedure involves identification of the “IT-preferred” layer(s) via a standard linear mapping to temporally averaged IT responses. *Choosing a temporal mapping gradation*. These “IT-preferred” model layer(s) are then mapped to 10*ms* timebins from 70-260*ms* in either a uniform or graded fashion, if the model is feedforward. For ConvRNNs, this temporal mapping is always one-to-one with these 10*ms* timebins. **(b) Defining model OSTs.** Once the temporal mapping has been defined, we train a linear SVM at each 10*ms* model timebin and compute the classifier’s *d*′ (displayed in each of the black dots for a given example image). The first timebin at which the model *d*′ matches the primate’s accuracy is defined as the “Model OST” for that image (obtained via linear interpolation), which is the same procedure previously used^12^ to determine the ground truth IT OST (“Primate OST” vertical dotted line). **(c) Proper choices of recurrence best match IT OSTs.** Mean and s.e.m. are computed across train/test splits (*N* = 10) when that image (of 1320 images) was a test-set image, with the Spearman correlation computed with the IT object solution times (analogously computed from the IT population responses) across the imageset solved by both the given model and IT, constituting the “Fraction of IT Solved Images” on the *x*-axis. We start with either a shallow base feedforward model consisting of 5 convolutional layers and 1 layer of readout (“BaseNet” in blue) as well as an intermediate-depth variant with 10 feedforward layers and 1 layer of readout (“BaseNet” in purple), detailed in Section A.2.1. From these base feedforward models, we embed recurrent circuits, resulting in either “Shallow ConvRNNs” or “Intermediate ConvRNNs”, respectively.

With model OST defined across both model families, we compared various ConvRNNs and feedforward models to the IT population’s OST in Figure 3c. Among shallower and deeper models, we found that ConvRNNs were generally able to better explain IT’s OST than their feedforward counterparts. Specifically, we found that ConvRNN circuits without *any* multiunit interaction such as the Time Decay ConvRNN only marginally, and not always significantly, improved the OST consistency over its respective BaseNet model^*b*^. On the other hand, ConvRNNs with multi-unit interactions generally provided the greatest match to IT OSTs than even the deepest feedforward models^*c*^, where the best feedforward model (ResNet-152) attains a mean OST consistency of 0.173 and the best ConvRNN (UGRNN) attains an OST consistency of 0.237.

Consistent with our observations in Figure 2 that different recurrent circuits with multi-unit interactions were not all equally effective when embedded in CNNs (despite outperforming the simple Time Decay model), we similarly found that this observation held for the case of matching IT’s OST. Given recent observations^32^ that inactivating parts of macaque ventrolateral PFC (vlPFC) results in behavioral deficits in IT for late-solved images, we reasoned that additional decoding procedures employed at the categorization layer during task optimization might meaningfully impact the model’s OST consistency, in addition to the choice of recurrent circuit used. We designed several decoding procedures (defined in Section A.5), motivated by prior observations of accumulation of relevant sensory signals during decision making in primates^33^. Overall, we found that ConvRNNs with different decoding procedures, but with the *same* layer-local recurrent circuit (RGC Median) and long-range feedback connectivity patterns, yielded significant differences in final consistency with the IT population OST (Figure S4; Friedman test, *p* < 0.05). Moreover, the simplest decoding procedure of outputting a prediction at the last timepoint, a strategy commonly employed by the computer vision community, had a lower OST consistency than each of the more nuanced Max Confidence^*d*^ and Threshold decoding procedures^*e*^ that we considered. Taken together, our results suggest that the type of multi-unit layer-wise recurrence *and* downstream decoding strategy are important features for OST consistency with IT, suggesting that specific, non-trivial connectivity patterns further downstream of the ventral visual pathway may be important to core object recognition behavior over timescales of a couple hundred milliseconds.

### 2.3 Neural dynamics differentiate ConvRNN circuits

ConvRNNs naturally produce a dynamic time series of outputs given an unchanging input stream, unlike feedforward networks. While these recurrent dynamics could be used for tasks involving time, here we optimized the ConvRNNs to perform the “static” task of object classification on ImageNet. It is possible that the primate visual system is optimized for such a task, because even static images produce reliably dynamic neural response trajectories at temporal resolutions of tens of milliseconds^15^. The object content of some images becomes decodable from the neural population significantly later than the content of other images, even though animals recognize both object sets equally well. Interestingly, late-decoding images are not well characterized by feedforward CNNs, raising the possibility that they are encoded in animals through recurrent computations^12^. If this were the case, we reason then that recurrent networks trained to perform a difficult, but static object recognition task might explain neural *dynamics* in the primate visual system, just as feedforward models explain time-averaged responses^34,35^.

Prior studies^23^ have *directly* fit recurrent parameters to neural data, as opposed to optimizing them on a task. While it is natural to try to fit recurrent parameters to the temporally-varying neural responses directly, this approach naturally has a loss of normative explanatory power. In fact, we found that this approach suffers from a fundamental overfitting issue to the particular image statistics of the neural data collected. Specifically, we directly fit these recurrent parameters (implanted into the task-optimized feedforward BaseNet) to the dynamic firing rates of primate neurons recorded during encoding of visual stimuli. However, while these non-task optimized dynamics generalized to held-out images and neurons (Figure S5a,b), they had no longer retained performance to the original object recognition task that the primate itself is able to perform (Figure S5c). Therefore, to avoid this issue, we instead asked whether *fully* task-optimized ConvRNN models (including the ones introduced in Section 2.1) could predict these dynamic firing rates from multi-electrode array recordings from the ventral visual pathway of rhesus macaques^36^.

We began with the feedforward BaseNet and added a variety of ConvRNN circuits, including the RGC Median ConvRNN and its counterpart generated at the random phase of the evolutionary search (“RGC Random”). All of the ConvRNNs were presented with the same images shown to the primates, and we collected the time series of features from each model layer. To decide which layer should be used to predict which neural responses, we fit linear models from each feedforward layer’s features to the neural population and measured where explained variance on held-out images peaked (see Section A.6 for more details). Units recorded from distinct arrays – placed in the successive V4, posterior IT (pIT), and central/anterior IT (cIT/aIT) cortical areas of the macaque – were fit best by the successive layers of the feedforward model, respectively. Finally, we measured how well ConvRNN features from these layers predicted the dynamics of each unit. In contrast with feedforward models fit to temporally-averaged neural responses, the linear mapping in the temporal setting must be *fixed* at all timepoints. The reason for this choice is that the linear mapping yields “artificial units” whose activity can change over time (just like the real target neurons), but the identity of these units should not change over the course of 260*ms*, which would be the case if instead a separate linear mapping was fit at each 10*ms* timebin. This choice of a temporally-fixed linear mapping therefore maintains the physical relationship between real neurons and model neurons.

As can be seen from Figure 4a, with the exception of the RGC Random ConvRNN, the ConvRNN feature dynamics fit the neural response trajectories as well as the feedforward baseline features on early phase responses (Wilcoxon test *p*-values in Table 1) and better than the feedforward baseline features for late phase responses (Wilcoxon test with Bonferroni correction *p* < 0.001), across V4, pIT, and cIT/aIT on held-out images. For the early phase responses, the ConvRNNs that employ direct passthrough are elaborations of the baseline feedforward network, although the ConvRNNs which only employ gating are still a nonlinear function of their input, similar to a feedforward network. For the late phase responses, any feedforward model exhibits similar “square wave” dynamics as its 100*ms* visual input, making it a poor predictor of the subset of late responses that are beyond the initial feedforward pass (Figure S6, purple lines). In contrast, the activations of ConvRNN units have persistent dynamics, yielding predictions of the *entire* neural response trajectories.

**Figure 4:**
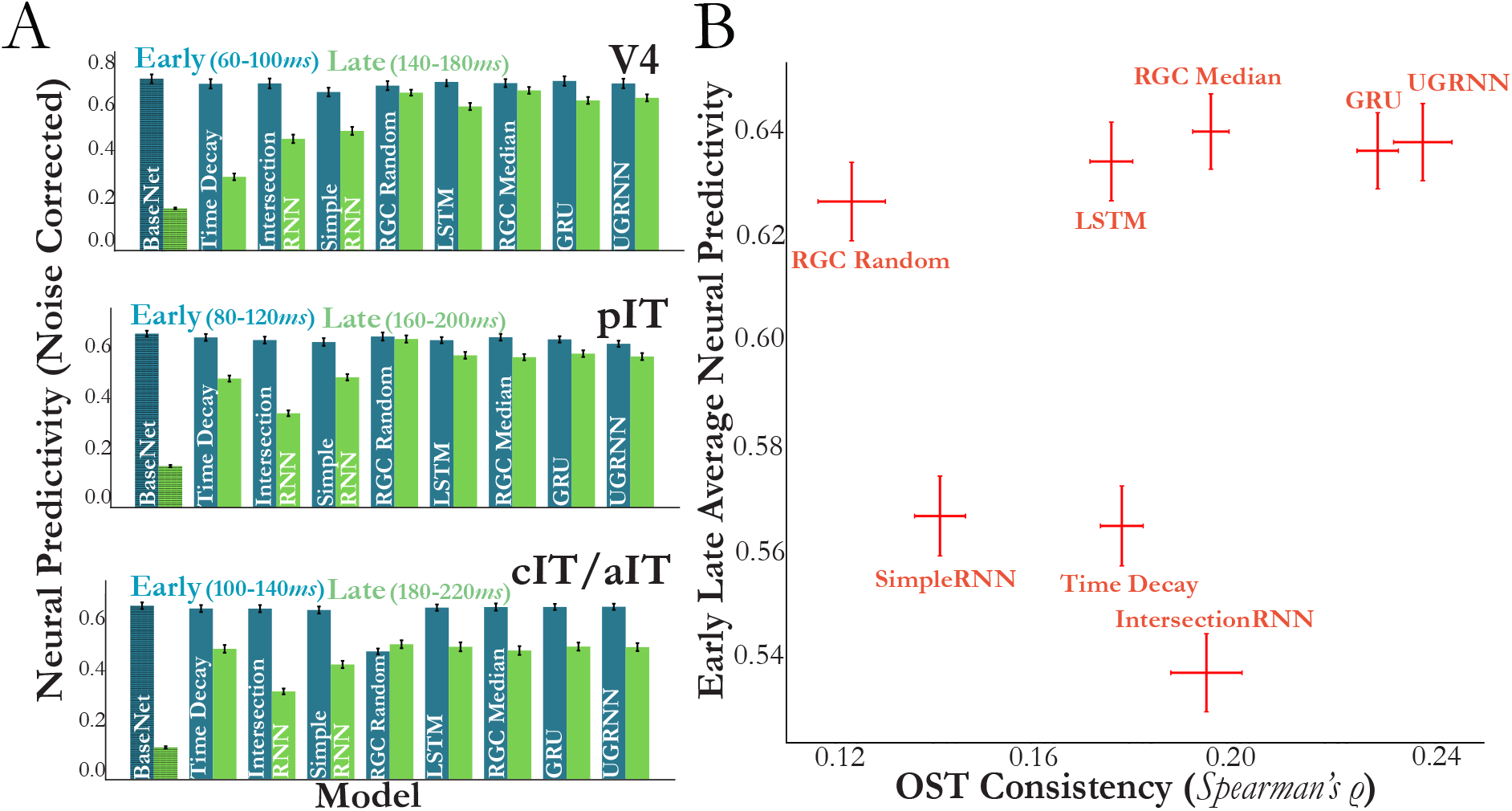
Suitably-chosen intermediate ConvRNN circuits provide consistent predictions of primate ventral stream neural dynamics. **(a)** The *y*-axis indicates the median across neurons of the explained variance between predictions and ground-truth responses on held-out images divided by the square root of the internal consistencies of the neurons, defined in Section A.6.3. Error bars indicates the s.e.m across neurons (*N* = 88 for V4, *N* = 88 for pIT, *N* = 80 for cIT/aIT) averaged across 10*ms* timebins (*N* = 4 each for the “Early” and “Late” designations). As can be seen, the intermediate-depth feedforward BaseNet model (first bars) is a poor predictor of the subset of late responses that are beyond the feedforward pass, but certain types of ConvRNN circuits (such as “RGC Median”, “UGRNN”, and “GRU”) added to the BaseNet are overall best predictive across visual areas at late timepoints (Wilcoxon test (with Bonferroni correction) with feedforward BaseNet, *p* < 0.001 for each visual area). See Figure S6 for the full timecourses at the resolution of 10*ms* bins. **(b)** For each ConvRNN circuit, we compare the average neural predictivity (averaged per neuron across early and late timepoints) averaged across areas, to the OST consistency. The ConvRNNs that have the best average neural predictivity also best match the OST consistency (“RGC Median”, “UGRNN”, and “GRU”).

**Table 1:** Wilcoxon test (with Bonferroni correction) *p*-values for comparing each intermediate-depth ConvRNN’s neural predictivity at the “early” timepoints (Figure 4) to the (11-layer) BaseNet.

| Model | Visual Area | Wilcoxon test $p$-value |
| --- | --- | --- |
| Time Decay | V4 | $< 0.001$ |
| IntersectionRNN | V4 | $< 0.001$ |
| LSTM | V4 | $< 0.001$ |
| UGRNN | V4 | $< 0.001$ |
| GRU | V4 | $< 0.001$ |
| SimpleRNN | V4 | $< 0.001$ |
| RGC Random | V4 | $< 0.001$ |
| RGC Median | V4 | $< 0.01$ |
| Time Decay | pIT | 0.022 |
| IntersectionRNN | pIT | $< 0.001$ |
| LSTM | pIT | $< 0.001$ |
| UGRNN | pIT | $< 0.001$ |
| GRU | pIT | $< 0.001$ |
| SimpleRNN | pIT | $< 0.001$ |
| RGC Random | pIT | 0.31 |
| RGC Median | pIT | $< 0.001$ |
| Time Decay | aIT | $< 0.001$ |
| IntersectionRNN | aIT | $< 0.001$ |
| LSTM | aIT | 0.47 |
| UGRNN | aIT | 0.09 |
| GRU | aIT | 0.16 |
| SimpleRNN | aIT | $< 0.001$ |
| RGC Random | aIT | $< 0.001$ |
| RGC Median | aIT | $< 0.01$ |

Crucially, these predictions result from the task-optimized nonlinear dynamics from ImageNet, as both models are fit to neural data with the same form of temporally-fixed linear model with the *same* number of parameters. Since the initial phase of neural dynamics was well-fit by feedforward models, we asked whether the later phase could be fit by a much simpler model than any of the ConvRNNs we considered, namely the Time Decay ConvRNN with ImageNet-trained time constants at convolutional layers. If the Time Decay ConvRNN were to explain neural data as well as the other ConvRNNs, it would imply that interneuronal recurrent connections are not needed to account for the observed dynamics; however, this model did not fit the late phase dynamics of intermediate areas (V4 and pIT) as well as the other ConvRNNs^*f*^. The Time Decay model did match the other ConvRNNs for cIT/aIT, which may indicate some functional differences in the temporal processing of this area versus V4 and pIT. Thus, the more complex recurrence found in ConvRNNs is generally needed both to improve object recognition performance over feedforward models *and* to account for neural dynamics in the ventral stream, even when animals are only required to fixate on visual stimuli. However, not all forms of complex recurrence are equally predictive of temporal dynamics. As depicted in Figure 4b, we found among these that the RGC Median, UGRNN, and GRU ConvRNNs attained the highest median neural predictivity for each visual area in both early and late phases, but in particular significantly outperformed the SimpleRNN ConvRNN at the late phase dynamics of these areas^*g*^, and these models in turn were among the best matches to IT object solution times (OST) from Section 2.2.

A natural follow-up question to ask is whether a *lack* of recurrent processing is the reason for the prior observation that there is a drop in explained variance for feedforward models from early to late timebins^12^. In short, we find that this is not the case, and that this drop likely has to do with task-orthogonal dynamics specific to individual primates, which we examine below.

It is well-known that recurrent neural networks can be viewed as very deep feedforward networks with weight sharing across layers that would otherwise be recurrently connected^16^. Thus, to address this question, we compare feedforward models of varying depths to ConvRNNs across the entire temporal trajectory under a *varying* linear mapping at each timebin, in contrast to the above. This choice of linear mapping allows us to evaluate how well the model features are at explaining early compared to late time dynamics without information from the early dynamics influencing the later dynamics, and also more crucially, to allow the feedforward model features to be independently compared to the late dynamics. Specifically, as can be seen in Figure S7a, we observe a drop in explained variance from early (130-140*ms*) to late (200-210*ms*) timebins for the BaseNet and ResNet-18 models, across multiple neural datasets. Models with increased feedforward depth (such as ResNet-101 or ResNet-152), along with our performance-optimized RGC Median ConvRNN, exhibit a similar drop in median population explained variance as the intermediate feedforward models. The benefit of model depth with respect to increased explained variance of late IT responses might be only noticeable while comparing shallow models (< 7 nonlinear transforms) to much deeper (> 15 nonlinear transforms) models^12^. Our results suggest that the amount of variance explained in the late IT responses is not a monotonically increasing function of model depth.

As a result, an alternative hypothesis is that the drop in explained variance from early to late timebins could instead be attributed to task-orthogonal dynamics specific to an individual primate as opposed to iterated nonlinear transforms, resulting in variability unable to be captured by any task-optimized model (feedforward or recurrent). To explore this possibility, we examined whether the model’s neural predictivity at these early and late timebins was relatively similar in ratio to that of one primate’s IT neurons mapped to that of another primate (see Section A.7 for more details, where we derive a novel measure of the the neural predictivity between animals, known as the “inter-animal consistency”).

As shown in Figure S7b, across various hyperparameters of the linear mapping, we observe a ratio close to one between the neural predictivity (of the target primate neurons) of the feedforward BaseNet to that of the source primate mapped to the same target primate. Therefore, as it stands, *temporally-varying* linear mappings to neural responses collected from an animal during rapid visual stimulus presentation (RSVP) may not sufficiently separate feedforward models from recurrent models any better than one animal to another – though more investigation is needed to ensure tight estimates of the inter-animal consistency measure we have introduced here with neural data recorded from more primates. Nonetheless, this observation further motivates our earlier result of additionally turning to temporally-varying *behavioral* metrics (such as the OST consistency), in order to be able to separate these model classes beyond what is currently achievable by neural response predictions.

### 2.4 ConvRNNs mediate a tradeoff between task performance and network size

Why might a suitably shallower feedforward network with temporal dynamics be desirable for the ventral visual stream? We reasoned that recurrence mediates a tradeoff between network size and task performance; a tradeoff that the ventral stream also maintains. To examine this possibility, in Figure 5, we compared each network’s task performance versus its size, measured either by parameter count or unit count. Across models, we found unit count (related to the number of neurons) to be more consistent with task performance than parameter count (related to the number of synapses). In fact, there are many models with a large parameter count but not very good task performance, indicating that adding synapses is not necessarily as useful for performance as adding neurons. For shallow recurrent networks, task performance seemed to be more strongly associated with OST consistency than network size. This tradeoff became more salient for deeper feedforward models and the intermediate ConvRNNs, as the very deep ResNets (ResNet-34 and deeper) attained an overall *lower* OST consistency compared to the intermediate ConvRNNs, using both much more units and parameters compared to small relative gains in task performance. Similarly, intermediate ConvRNNs with high task performance and minimal *unit* count, such as the UGRNN, GRU, and RGCs attained both the highest OST consistency overall (Figures 3 and 5) along with providing the best match to neural dynamics among ConvRNN circuits across visual areas (Figure 4b). This observation indicates that suitably-chosen recurrence can provide a means for maintaining this fundamental tradeoff.

**Figure 5:**
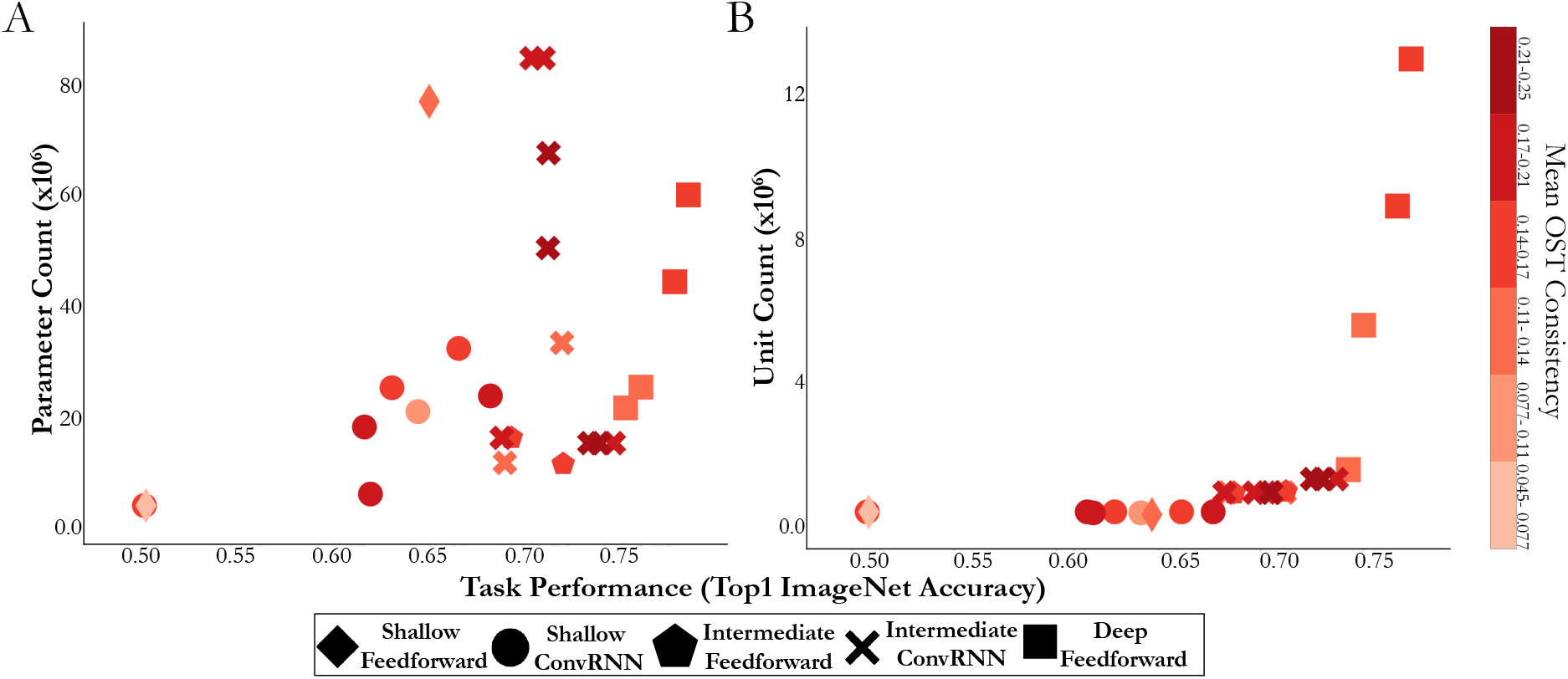
Intermediate ConvRNN circuits with highest OST consistency conserve on network size while maintaining task performance. Across all models considered, the intermediate ConvRNNs (denoted by “x”) that attain high categorization performance (*x*-axis) while maintaining a low unit count (panel B) rather than parameter count (panel A) for their given performance level, achieve the highest mean OST consistency (Spearman correlation with IT population OST, averaged across *N* = 10 train/test splits). The colorbar indicates this mean OST consistency (monotonically increasing from purple to red), binned into 6 equal ranges. Models with a larger network size at a fixed performance level are *less consistent* with primate object recognition behavior (e.g. deep feedforward models, denoted by boxes), with recurrence maintaining a fundamental tradeoff between network size and task performance.

Given our finding that specific forms of task-optimized recurrence are more consistent with IT’s OST than iterated feedforward transformations (with unshared weights), we asked whether it was possible to approximate the effect of recurrence with a feedforward model. This approximation would allow us to better describe the additional “action” that recurrence is providing in its improved OST consistency. In fact, one difference between this metric and the explained variance metric evaluated on neural responses in the prior section is that the latter uses a linear transform from model features to neural responses, whereas the former operates directly on the original model features. Therefore, a related question is whether the (now standard) use of a linear transform for mapping from model units to neural responses can potentially *mask* the behavioral improvement that suitable recurrent processing has over deep feedforward models in their original feature space.

To address these questions, we trained a separate linear mapping (PLS regression) from each model layer to the corresponding IT response at the given timepoint, on a set of images distinct from those on which the OST consistency metric is evaluated on (see Section A.8.2 for more details). The outputs of this linear mapping were then used in place of the original model features for both the uniform and graded mappings, constituting “PLS Uniform” and “PLS Graded”, respectively. Overall, as depicted in Figure S3, we found that models with *less* temporal variation in their source features (namely, those under a uniform mapping with less “IT-preferred” layers than the total number of timebins) had significantly *improved* OST consistency with their linearly transformed features under PLS regression (Wilcoxon test, *p* < 0.001; mean OST difference 0.0458 and s.e.m. 0.00399). On the other hand, the linearly transformed intermediate feedforward models were *not* significantly different from task-optimized ConvRNNs that achieved high OST consistency^*h*^, suggesting that the action of suitable task-optimized recurrence approximates that of a shallower feedforward model with linearly induced ground-truth neural dynamics.

## Discussion

The overall goal of this study is to determine what role recurrent circuits may have in the execution of core object recognition behavior in the ventral stream. By broadening the method of goal-driven modeling from solving tasks with feedforward CNNs to ConvRNNs that include layer-local recurrence and feedback connections, we first demonstrate that appropriate choices of these recurrent circuits which incorporate specific mechanisms of “direct passthrough” and “gating” lead to matching the task performance of much deeper feedforward CNNs with fewer units and parameters, even when minimally unrolled. This observation suggests that the recurrent circuit motif plays an important role even during the initial timepoints of visual processing. Moreover, unlike very deep feedforward CNNs, the mapping from the early, intermediate, and higher layers of these ConvRNNs to corresponding cortical areas is neuroanatomically consistent and reproduces prior quantitative properties of the ventral stream. In fact, ConvRNNs with high task performance but small network size (as measured by number of neurons rather than synapses) are most consistent with the temporal evolution of primate IT object identity solutions. We further find that these task-optimized ConvRNNs can reliably produce quantitatively accurate dynamic neural response trajectories at temporal resolutions of tens of milliseconds throughout the ventral visual hierarchy.

Taken together, our results suggest that recurrence in the ventral stream extends feedforward computations by mediating a tradeoff between task performance and neuron count during core object recognition, suggesting that the computer vision community’s solution of stacking more feedforward layers to solve challenging visual recognition problems approximates what is compactly implemented in the primate visual system by leveraging additional nonlinear temporal transformations to the initial feedforward IT response. This work therefore provides a quantitative prescription for the next generation of dynamic ventral stream models, addressing the call to action in a recent previous study^12^ for a change in architecture from feedforward models.

Many hypotheses about the role of recurrence in vision have been put forward, particularly in regards to overcoming certain challenging image properties^5,6,7,8,4,9,10,11,12,13,14,15^. We believe this is the first work to address the role of recurrence at scale by connecting novel *task-optimized* recurrent models to temporal metrics defined on high-throughput neural and behavioral data, to provide evidence for recurrent connections extending feedforward computations. Moreover, these metrics are well-defined for feedforward models (unlike prior work^19^) and therefore meaningfully demonstrate a separation between the two model classes.

Though our results help to clarify the role of recurrence during core object recognition behavior, many major questions remain. Our work addresses why the visual system may leverage recurrence to subserve visually challenging behaviors, replacing a physically implausible architecture (deep feedforward CNNs) with one that is ubiquitously consistent with anatomical observations (ConvRNNs). However, our work does not address gaps in understanding either the loss function or the learning rule of the neural network. Specifically, we intentionally implant layer-local recurrence and long-range feedback connections into feedforward networks that have been useful for supervised learning via backpropagation on ImageNet. A natural next step would be to connect these ConvRNNs with unsupervised objectives, as has been done for feedforward models of the ventral stream in concurrent work^37^. The question of biologically plausible learning targets is similarly linked to biologically plausible mechanisms for learning such objective functions. Recurrence could play a separate role in implementing the propagation of error-driven learning, obviating the need for some of the issues with backpropagation (such as weight transport), as has been recently demonstrated at scale^38,39^. Therefore, building ConvRNNs with unsupervised objective functions optimized with biologically-plausible learning rules would be essential towards a more complete goal-driven theory of visual cortex.

Additionally, high-throughput experimental data will also be critical to further separate hypotheses about recurrence. While we see evidence of recurrence as mediating a tradeoff between network size and task performance for core object recognition, it could be that recurrence plays a more task-specific role under more temporally dynamic behaviors. Not only would it be an interesting direction to optimize ConvRNNs on more temporally dynamic visual tasks than ImageNet, but to compare to neural and behavioral data collected from such stimuli, potentially over longer timescales than 260*ms*. While the RGC motif of gating and direct passthrough gave the highest task performance among ConvRNN circuits, the circuits that maintain a tradeoff between number of units and task performance (RGC Median, GRU, and UGRNN) had the best match to the current set of neural and behavioral metrics, even if some of them do not employ passthroughs. However, it could be the case that with the same metrics we develop here but used in concert with such stimuli over potentially longer timescales, that we can better differentiate these three ConvRNN circuits. Therefore, such models and experimental data would synergistically provide great insight into how rich visual behaviors proceed, while also inspiring better computer vision algorithms.

## Acknowledgements

We thank Tyler Bonnen, Eshed Margalit, and the anonymous reviewers for comments on this manuscript. We thank the Google TensorFlow Research Cloud (TFRC) team for generously providing TPU hardware resources for this project. D.L.K.Y is supported by the James S. McDonnell Foundation (Understanding Human Cognition Award Grant No. 220020469), the Simons Foundation (Collaboration on the Global Brain Grant No. 543061), the Sloan Foundation (Fellowship FG-2018-10963), the National Science Foundation (RI 1703161 and CAREER Award 1844724), the DARPA Machine Common Sense program, and hardware donation from the NVIDIA Corporation. This work is also supported in part by Simons Foundation grant SCGB-542965 (J.J.D. & D.L.K.Y.). This project has received funding from the European Union’s Horizon 2020 research and innovation programme under grant agreement No. 70549 (J.K.). J.S. is supported by the Mexican National Council of Science and Technology (CONACYT).

## Author Contributions

A.N. and D.L.K.Y. designed the experiments. A.N., J.S., and D.B. conducted the experiments, and A.N. analyzed the data. K.K. contributed neural data, and J.K. contributed to initial code development. K.K. and J.J.D. provided technical advice on neural predictivity metrics. D.S. and S.G. provided technical advice on recurrent neural network training. A.N. and D.L.K.Y. interpreted the data and wrote the paper.

## Competing Interest Declaration

The authors declare no competing interests.

## A Methods

### A.1 Model framework

#### A.1.1 Software package

To explore the architectural space of ConvRNNs and compare these models with the primate visual system, we used the Tensorflow library^40^ to augment standard CNNs with both local and long-range recurrence (Figure 1). Conduction from one area to another in visual cortex takes approximately 10*ms*^41^, with signal from photoreceptors reaching IT cortex at the top of the ventral stream by 70-100*ms*. Neural dynamics indicating potential recurrent connections take place over the course of 100-260*ms*^15^. A single feedforward volley of responses thus cannot be treated as if it were instantaneous relative to the timescale of recurrence and feedback. Hence, rather than treating each entire feedforward pass from input to output as one integral time step, as is normally done with RNNs^5^, each time step in our models corresponds to a single feedforward layer processing its input and passing it to the next layer. This choice required an unrolling scheme different from that used in the standard Tensorflow RNN library, the code for which, including trained model weights, can be found on our Github repository: https://github.com/neuroailab/convrnns.

#### A.1.2 Defining ConvRNNs

Within each ConvRNN layer, feedback inputs from higher layers are resized to match the spatial dimensions of the feedforward input to that layer. Both types of input are processed by standard 2D convolutions. If there is any local recurrence at that layer, the output is next passed to the recurrent circuit as input. Feedforward and feedback inputs are combined within the recurrent circuit by spatially resizing the feedback inputs (via bilinear interpolation) and concatenating these with the feedforward input across the channel dimension. We let ⊕ denote this concatenation along the channel dimension with appropriate resizing to align spatial dimensions. Finally, the output of the circuit is passed through any additional nonlinearities, such as max-pooling. The generic update rule for the discrete-time trajectory of such a network is thus:

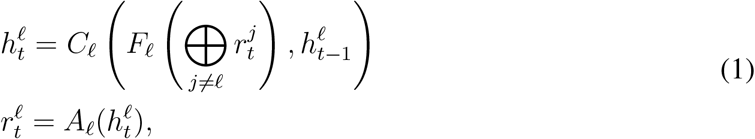

where 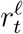 is the output of layer ℓ at time *t*, 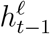 is the hidden state of the locally recurrent circuit *C*_ℓ_ at time *t* – 1, and *A*_ℓ_ is the activation function and any pooling post-memory operations. The learned parameters of such a network consist of *F*_ℓ_, comprising any feedforward and feedback connections coming into layer ℓ = 1, …, *L*, and any of the learned parameters associated with the local recurrent circuit *C*_ℓ_.

In this work, all forms of recurrence add parameters to the feedforward base model. Because this could improve task performance for reasons unrelated to recurrent computation, we trained two types of control model to compare to ConvRNNs:

1. Feedforward models with more convolution filters (“wider”) or more layers (“deeper”) to approximately match the number of parameters in a recurrent model.
2. Replicas of each ConvRNN model unrolled for a *minimal* number of time steps, defined as the number that allows all model parameters to be used at least once. A minimally unrolled model has exactly the same number of parameters as its fully unrolled counterpart, so any increase in performance from unrolling longer can be attributed to recurrent computation. Fully and minimally unrolled ConvRNNs were trained with identical learning hyperparameters.

#### A.1.3 Training Procedure

All models (both feedforward and ConvRNN) used the standard ResNet preprocessing provided by TensorFlow here: https://github.com/tensorflow/tpu/blob/master/models/official/resnet/resnet_preprocessing.py. Furthermore, they were trained on 224 pixel ImageNet with stochastic gradient descent with momentum (SGDM)^42^, using a momentum value of 0.9.

We allowed the base learning rate, batch size, and L2 regularization strength to vary for each model, depending on what was optimal in terms of top-1 validation accuracy for that model. All models (except for AlexNet) used the ResNet training schedule^24^, whereby the base learning rate is decayed by 90% at 30, 60, and 80 epochs, training for 90 epochs total. The AlexNet had its base learning rate of 0.01 subsequently decayed to 0.005, 0.001, and 0.0005, at 30, 60, and 80 epochs, respectively. We list these values for each model in the table below:

| Model Class | Base Learning Rate | Batch Size | L2 Regularization |
| --- | --- | --- | --- |
| AlexNet | 0.01 | 1024 | $5 \times 10^{-4}$ |
| 6-layer BaseNet | 0.01 | 256 | $1 \times 10^{-4}$ |
| Shallow ConvRNNs | 0.01 | 256 | $1 \times 10^{-4}$ |
| 11-layer BaseNet | 0.0025 | 64 | $1 \times 10^{-4}$ |
| ResNets | 0.025 | 64 | $1 \times 10^{-4}$ |
| Intermediate ConvRNNs | 0.0025 | 64 | $1 \times 10^{-4}$ |

The only exceptions to the above are the models that are the result of the large-scale hyperparameter searches, detailed in Section A.4. Here the learning rate and batch size are allowed to vary, and the L2 regularization is not uniform across the model, but is also allowed to vary for both the feedforward backbone and each layer’s ConvRNN circuit. We list the learning rates and batch sizes for these models below:

| Model | Base Learning Rate | Batch Size |
| --- | --- | --- |
| Shallow LSTM (“LSTM Opt” in Figure S1) | $7.587 \times 10^{-3}$ | 64 |
| RGC Random | $5.184 \times 10^{-3}$ | 64 |
| RGC Median | $6.736 \times 10^{-3}$ | 64 |

Since these model hyperparameters are non-standard, we manually drop the learning rate (using the same decay factor of 90%) once the top-1 validation accuracy saturates at that given learning rate.

### A.2 Feedforward model architectures

#### A.2.1 BaseNet architectures

Here we provide the architectures of the feedforward CNNs we developed in this paper, referred to as “BaseNet” when they are later implanted with ConvRNN circuits. For all of these architectures, we use ELU nonlinearities^43^.

The 6-layer BaseNet (into which we implanted ConvRNN circuits to form the orange “Shallow ConvRNN” model class in Figure 3c), referenced as “FF” in Figure S1, referred to as “BaseNet” among the blue “Shallow Feedforward” models in Figure 3c, and “Feedforward” in Figure S5c, had the following architecture:

| Layer | Kernel Size | Channels | Stride | Max Pooling |
| --- | --- | --- | --- | --- |
| 1 | $7 \times 7$ | 64 | 2 | $2 \times 2$ |
| 2 | $3 \times 3$ | 128 | 1 | $2 \times 2$ |
| 3 | $3 \times 3$ | 256 | 1 | $2 \times 2$ |
| 4 | $3 \times 3$ | 256 | 1 | $2 \times 2$ |
| 5 | $3 \times 3$ | 512 | 1 | $2 \times 2$ |
| 6 | $2 \times 2$ | 1000 | 1 | No |

The 11-layer BaseNet used for the “Intermediate ConvRNNs” (red models in Figure 3c) and modeled after ResNet-18^24^ (but using MaxPooling rather than stride-2 convolutions to perform downsampling) is given below:

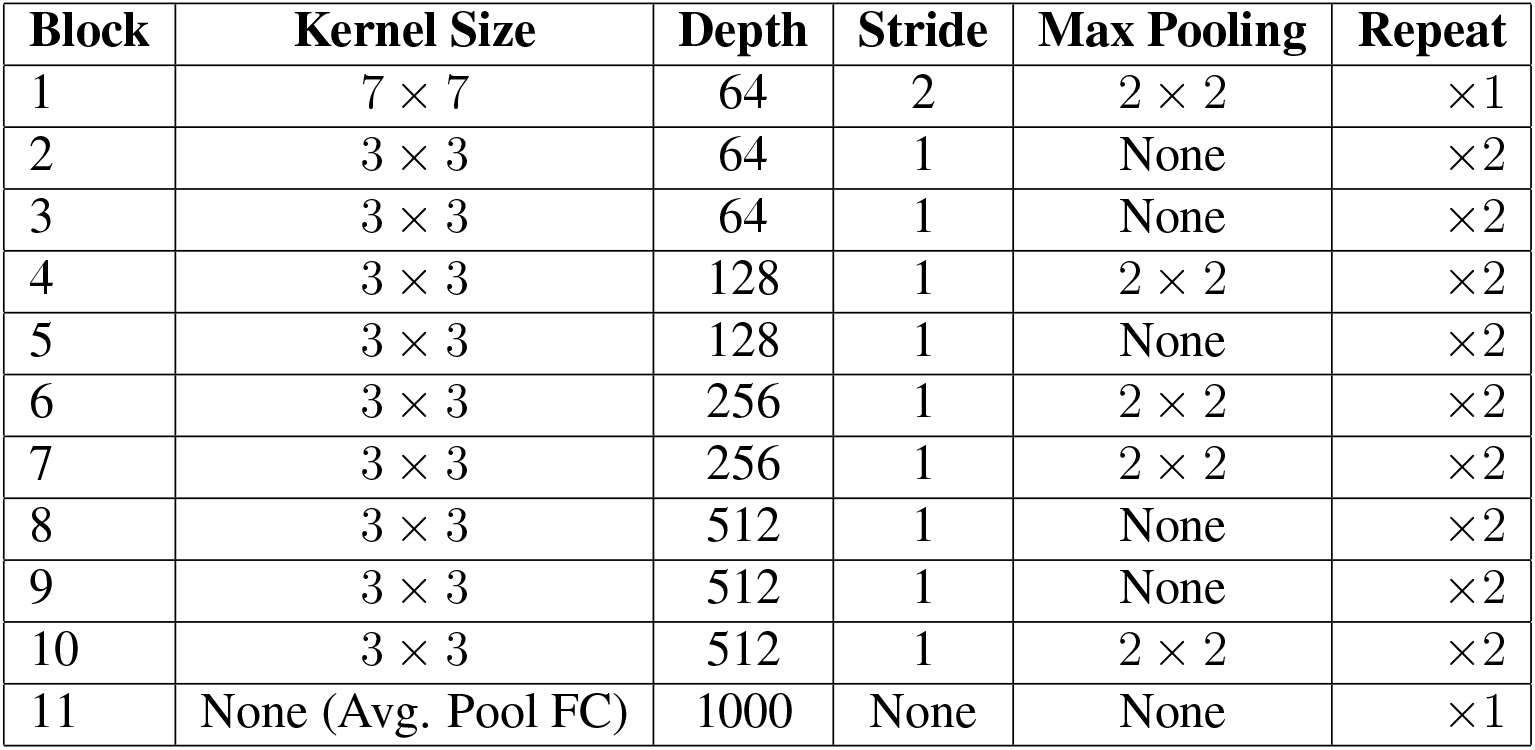

| Block | Kernel Size | Depth | Stride | Max Pooling | Repeat |
| --- | --- | --- | --- | --- | --- |
| 1 | $7 \times 7$ | 64 | 2 | $2 \times 2$ | $\times 1$ |
| 2 | $3 \times 3$ | 64 | 1 | None | $\times 2$ |
| 3 | $3 \times 3$ | 64 | 1 | None | $\times 2$ |
| 4 | $3 \times 3$ | 128 | 1 | $2 \times 2$ | $\times 2$ |
| 5 | $3 \times 3$ | 128 | 1 | None | $\times 2$ |
| 6 | $3 \times 3$ | 256 | 1 | $2 \times 2$ | $\times 2$ |
| 7 | $3 \times 3$ | 256 | 1 | $2 \times 2$ | $\times 2$ |
| 8 | $3 \times 3$ | 512 | 1 | None | $\times 2$ |
| 9 | $3 \times 3$ | 512 | 1 | None | $\times 2$ |
| 10 | $3 \times 3$ | 512 | 1 | $2 \times 2$ | $\times 2$ |
| 11 | None (Avg. Pool FC) | 1000 | None | None | $\times 1$ |

This is the BaseNet among the purple “Intermediate Feedforward” models in Figure 3c, and used in Figures 4, S3, S6, and S7.

The variant of the above 6-layer feedforward CNN, referenced in Figure S1 as “FF Wider” is given below:

| Layer | Kernel Size | Channels | Stride | Max Pooling |
| --- | --- | --- | --- | --- |
| 1 | $7 \times 7$ | 128 | 2 | $2 \times 2$ |
| 2 | $3 \times 3$ | 512 | 1 | $2 \times 2$ |
| 3 | $3 \times 3$ | 512 | 1 | $2 \times 2$ |
| 4 | $3 \times 3$ | 512 | 1 | $2 \times 2$ |
| 5 | $3 \times 3$ | 1024 | 1 | $2 \times 2$ |
| 6 | $2 \times 2$ | 1000 | 1 | None |

The “FF Deeper” model referenced in Figure S1 is given below:

| Layer | Kernel Size | Depth | Stride | Max Pooling |
| --- | --- | --- | --- | --- |
| 1 | $7 \times 7$ | 64 | 2 | $2 \times 2$ |
| 2 | $3 \times 3$ | 64 | 1 | None |
| 3 | $3 \times 3$ | 64 | 1 | None |
| 4 | $3 \times 3$ | 128 | 1 | $2 \times 2$ |
| 5 | $3 \times 3$ | 128 | 1 | None |
| 6 | $3 \times 3$ | 256 | 1 | $2 \times 2$ |
| 7 | $3 \times 3$ | 256 | 1 | $2 \times 2$ |
| 8 | $3 \times 3$ | 512 | 1 | None |
| 9 | $3 \times 3$ | 512 | 1 | None |
| 10 | $3 \times 3$ | 512 | 1 | $2 \times 2$ |
| 11 | None (Avg. Pool FC) | 1000 | None | None |

#### A.2.2 AlexNet

We use the standard AlexNet architecture, which uses local response normalization^26^. We note that we are able to attain a higher than reported top-1 validation accuracy of 63.9% (compared to 57% accuracy) by using the ResNet preprocessing mentioned in Section A.1.3.

#### A.2.3 ResNet Architectures

For the ResNet architectures, we used the original v1 versions^24^ for ResNet-18 and ResNet-34. For deeper ResNets (ResNet-50, ResNet-101, and ResNet-152), we used the v2 variants of ResNets, as this gave them a slightly higher increase in top-1 ImageNet validation accuracy. Specifically, the v2 variants of ResNets use the pre-activation of the weight layers rather than the post-activation used in the original versions. Furthermore, the v2 variants of ResNets apply batch normalization^44^ and ReLU to the input *prior* to the convolution, whereas the original variants apply these operations after the convolution. We use the TensorFlow Slim implementations for these two variants provided here: https://github.com/tensorflow/models/tree/master/research/slim.

### A.3 ConvRNN Circuit Equations

Here we provide the explicit update equations for each of the ConvRNN circuits referenced in the barplot in Figure 2c (*C*_ℓ_ in (1)), where *σ* denotes the sigmoid function.

Throughout these sections, we let ∘ denote Hadamard (elementwise) product, let * denote convolution, let 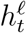 denote the output of the circuit, let 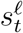 denote the propagated memory of the circuit (also known as the hidden state), and let 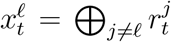 denote the input to the circuit at layer ℓ (this is the concatenation of feedforward and feedback inputs to layer ℓ, defined in Section A.1.2).

In the following table, we provide the number of timesteps the ConvRNNs were unrolled for during training (“Fully Unrolled”), what the corresponding minimally unrolled timesteps would be to engage recurrent connections once for each model class, and the number of timesteps for evaluation when comparing to neural and behavioral data:

| Model Class | Minimally Unrolled | Fully Unrolled | Evaluation |
| --- | --- | --- | --- |
| Shallow ConvRNNs | 7 | 16 | 26 |
| Intermediate ConvRNNs | 12 | 17 | 26 |
| RGC Random | 12 | 26 | 26 |

We also list the timestep at which the image presentation was replaced by a mean gray stimulus during model training and model evaluation:

| Model Class | Training | Evaluation |
| --- | --- | --- |
| Shallow ConvRNNs | 12 | 10 |
| Intermediate ConvRNNs | 12 | 10 |
| RGC Random | 10 | 10 |

The above training parameters were chosen based on what yielded high performance for that model class and also what was able to feasibly fit into TPU memory for training (more unroll timesteps requires more memory, but can also lead to instability during training, as is common with training RNNs^45^).

For the “Shallow ConvRNNs”, ConvRNN circuits were implanted into convolutional layers 3, 4, and 5 of the 6-layer BaseNet. For the “Intermediate ConvRNNs”, ConvRNN circuits were implanted into convolutional layers 4, 5, 6, 7, 8, 9, and 10 of the 11-layer BaseNet.

#### A.3.1 Time Decay

This is the simplest form of recurrence that we consider and has a discrete-time trajectory given by

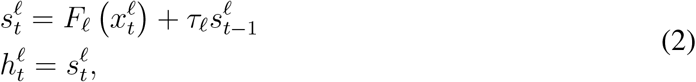

where *τ*_ℓ_ is the learned time constant at a given layer ℓ. This model is intended to be a control for simplicity, where the time constants could model synaptic facilitation and depression in a cortical layer.

For the TensorFlow implementation of this circuit, see the `GenFuncCell()` class in the `utils.cells.py` file on our Github repository.

#### A.3.2 SimpleRNN

The update equations in this case are given by:

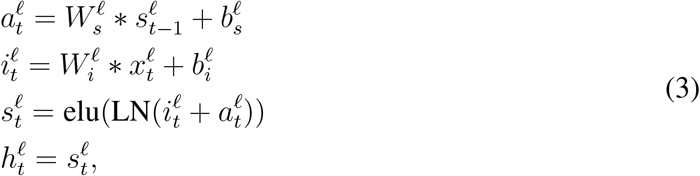

where LN denotes the layer normalization operation^46^ with offset parameter *β* initialized to 0 and scale parameter *γ* initialized to 1. For the shallow SimpleRNN (among the orange “Shallow ConvRNN” models in Figure 3c), we use layer normalization but omit its usage in the intermediate ConvRNN as it was not able to train with that operation.

For the TensorFlow implementation of this circuit, see the `ConvNormBasicCell()` class in the `utils.cells.py` file on our Github repository.

#### A.3.3 GRU

We adapt the standard GRU circuit^47^ to the convolutional setting:

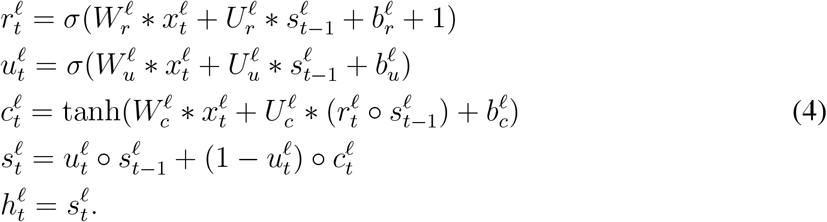

For the TensorFlow implementation of this circuit, see the `ConvGRUCell()` class in the `utils.cells.py` file on our Github repository.

#### A.3.4 LSTM

We adapt the standard LSTM circuit^48^ to the convolutional setting, with some slight modifications such as added layer normalization for stability in training.

We first make the gates convolutional as follows:

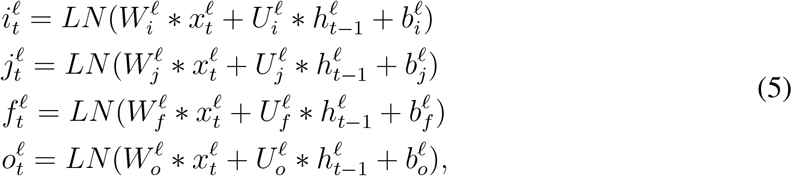

where LN denotes the layer normalization operation^46^ with offset parameter *β* initialized to 0 and scale parameter *γ* initialized to 1.

Next, the LSTM update equations are as follows:

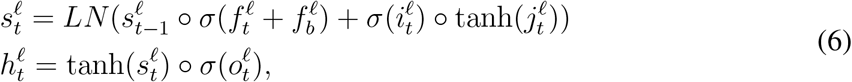

where 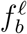 is the forget gate bias, typically set to 1, as recommended by others^49^. When peephole connections^50^ are allowed, these update equations are augmented to become:

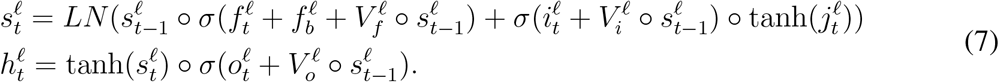

In the shallow LSTM (among the orange “Shallow ConvRNN” models in Figure 3c), we use peepholes and layer normalization, as that was found in the LSTM search for shallow models (described in Section A.4.1) to be useful for performance. We found, however, that neither of these augmentations are needed in the deeper variant (among the red “Intermediate ConvRNNs” in Figure 3c) in order to achieve high top-1 validation accuracy on ImageNet.

For the TensorFlow implementation of this circuit, see the `ConvLSTMCell()` class in the `utils.cells.py` file on our Github repository.

#### A.3.5 UGRNN

We adapt the UGRNN^30^ to the convolutional setting. The update equations are as follows:

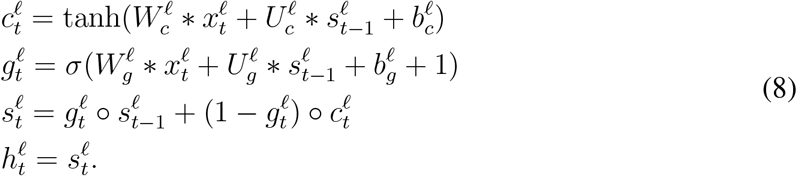

For the TensorFlow implementation of this circuit, see the `ConvUGRNNCell()` class in the `utils.cells.py` file on our Github repository.

#### A.3.6 IntersectionRNN

We adapt the IntersectionRNN^30^ to the convolutional setting. The update equations are as follows:

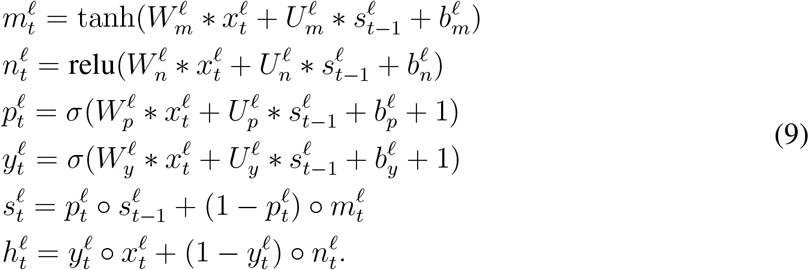

For the TensorFlow implementation of this circuit, see the `ConvIntersectionRNNCell()` class in the `utils.cells.py` file on our Github repository.

#### A.3.7 Reciprocal Gated Circuit (RGC)

Here we provide the explicit update equations for the Reciprocal Gated Circuit^29^, diagrammed in Figure 2a (bottom right). The update equation for the output of the circuit, 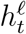, is given by a gating of both the input 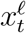 and prior output 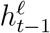:

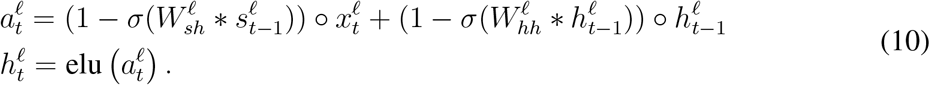

The update equation for the memory 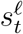 is given by a gating of the input 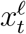 and the prior state 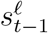:

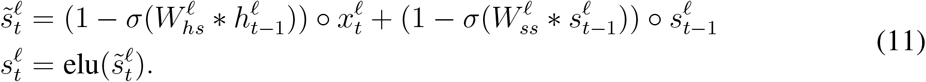

For the TensorFlow implementation of this circuit, see the `ReciprocalGateCell()` class in the `utils.cells.py` file on our Github repository.

### A.4 ConvRNN Searches

We employed a form of Bayesian optimization, a Tree-structured Parzen Estimator (TPE), to search the space of continuous and categorical hyperparameters^51^. This algorithm constructs a generative model of *P*[*score* | *configuration]* by updating a prior from a maintained history *H* of hyperparameter configuration-loss pairs. The fitness function that is optimized over models is the expected improvement, where a given configuration *c* is meant to optimize 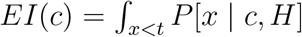. This choice of Bayesian optimization algorithm models *P*[*c* | *x*] via a Gaussian mixture, and restricts us to tree-structured configuration spaces.

Models were trained synchronously 100 models at a time using the HyperOpt package^52^, which implements the above Bayesian optimization. Each model was trained on its own Tensor Processing Unit (TPUv2), and during the search, ConvRNN models were trained by stochastic gradient descent on 128 pixel ImageNet for efficiency. The top performing ConvRNN models were then fully trained out on 224 pixel ImageNet.

#### A.4.1 LSTM search

The search for better LSTM architectures involved searching over training hyperparameters and common structural variants of the LSTM to better adapt this local structure to deep convolutional networks, using hundreds of second generation Google Tensor Processing Units (TPUv2s). We searched over learning hyperparameters (e.g. gradient clip values, learning rate) as well as structural hyperparameters (e.g. gate convolution filter sizes, channel depth, whether or not to use peephole connections, etc.).

Specifically, we implanted LSTMs into convolutional layers 3, 4, and 5, of the 6-layer BaseNet described in Section A.2. At each of these layers, the parameters of the LSTM circuit (defined in Section A.3.4) were allowed to vary per layer, as follows:

- The discrete number of convolutional channels was chosen from {64, 128, 256}.
- The discrete choice of convolutional filter sizes were chosen from {1, 4}.
- The binary choice of whether or not to use layer normalization.
- The strength of the L2 regularization of all LSTM parameters in that layer ∈ [10^−7^, 10^−3^], sampled log-uniformly.
- The scale of the He-style initialization^53^ of the convolutional filter weights ∈ [0.25, 2], sampled uniformly.
- The value of the constant initialization of the biases ∈ [–2, 2], sampled uniformly.
- The forget gate bias 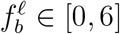, sampled uniformly (defined in (6)).
- The binary choice of whether or not to use peephole connections (as defined in (7)).

Outside of the LSTM circuit at each layer, we additionally searched over the following parameters as well:

- The number of discrete timesteps the model is unrolled ∈ [12, 26], sampled uniformly in consecutive groups of size 2.
- The timestep at each the image presentation is “turned off” and replaced with a mean gray stimulus ∈ [8, 12], sampled uniformly in consecutive groups of 2.
- The discrete choice of batch size used for the training the entire model ∈ {64, 128, 256}.
- The learning rate for training the entire model ∈ [10^−3^, 10^−1^], sampled log-uniformly.
- The binary choice of whether or not to use Nesterov momentum^54^.
- The gradient clipping value ∈ [0.3, 3], sampled log-uniformly.
- The scale of the He-style initialization^53^ of the convolutional filter weights of the feedforward base model ∈ [0.25, 2], sampled uniformly.
- The strength of the L2 regularization of the feedforward base model parameters ∈ [10^−7^, 10^−3^], sampled log-uniformly.

Each search point is a sampled value from the above described search space and trained for 1 epoch on ImageNet, in order to sample as many models as much as possible with the computational resources available. More than 1600 models were sampled in total, and we trained out the top ones and the median performing one after 1 epoch were trained out fully on 224 pixel ImageNet. The median model from this search attained the best top-1 validation accuracy on ImageNet, which is the resultant “LSTM Opt” model in Figure S1 and otherwise referred to as “Shallow LSTM”. The configuration of chosen hyperparameters for this model can be found in the `configs.lstm_shallow.npz` file on our Github repository.

#### A.4.2 Reciprocal Gated Circuit (RGC) search

From the Reciprocal Gated Circuit equations in (10) and (11), there are a variety of possibilities for how 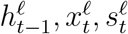, and 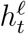 can be connected to one another (schematized in Figure 2b).

Mathematically, the search in Figure 2b can be formalized in terms of the following update equations. First, we define our input sets and building block functions:

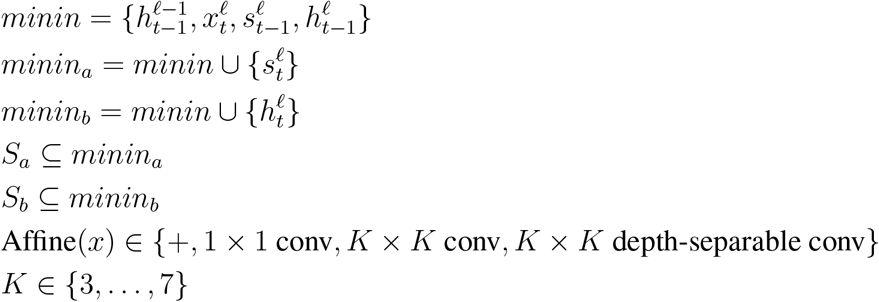

With those in hand, we have the following update equations:

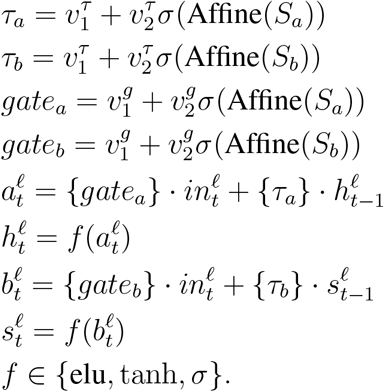

For clarity, the following matrix summarizes the connectivity possibilities (with ? denoting the possibility of a connection), schematized in Figure 2b:

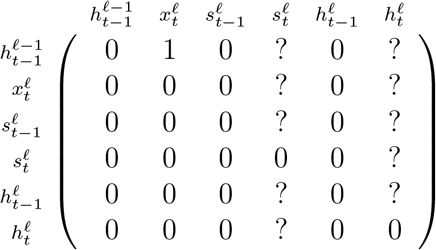

Each search point is a sampled value from the above described search space and trained for five epochs on ImageNet, in order to sample as many models as much as possible with the computational resources available. Around 6000 models were sampled in total over the course of the search. The top and median models from this search were then fully trained out on 224 pixel ImageNet with a batch size of 64 (which was maximum that we could fit into TPU memory). Moreover, as explicated in the table in Section A.1.3, the ResNet models were also trained using this same batch size, with the standard ResNet learning rate of 0.1 for a batch size of 256 linearly rescaled to accomodate, to ensure fair comparison between these two model classes. The median model from this search attained the best top-1 validation accuracy on ImageNet of all models selected to be trained out fully on ImageNet from the search, producing the resultant “RGC Median” model in Figure 2c (note that this designation also includes the long-range feedback connections). The configuration of chosen hyperparameters for this model can be found in the `configs.median_rgcell_cfg.py` file on our Github repository. The “RGC Random” model is from the random phase of this search (400th sampled model, since models sampled earlier than that failed to train out fully on ImageNet).

### A.5 Decoders

In addition to choice of ConvRNN circuit, we consider particular choices of “light-weight” (in terms of parameter count) decoding strategy that determines the final object category of that image. By construction, the model will output category logit probabilities at each timestep, given by the softmax function softmax 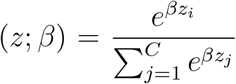, where *C* = 1000 is the number of ImageNet categories. This will then be passed to a decoding function which can take one of several forms:

1. **Default:** Use the logits at the last timestep and discard the remaining, with *β* =1.
2. **Threshold Decoder:** Select the logits from the first timepoint at which the maximum logit value at that timepoint crosses a fixed threshold (set to 0.9), with *β* =1.
3. **Max Confidence Decoder:** For the most confident category, find the timepoint at which that confidence peaks, and return the logits at that timepoint, where *β* is a trainable scalar parameter initialized to 1.

“RGC Median” therefore refers to the model trained using the default decoder, but when using the other two decoders with the “RGC Median” model, we append it to the name (as is done in Figures S3, S4, and S7a). The TensorFlow implementations of these decoders can be found in the `utils.decoders.py` file on our Github repository.

### A.6 Model prediction of neural responses

#### A.6.1 Neural data

Neural responses came from three multi-unit arrays per primate (rhesus macques): one implanted in V4, one in posterior IT (pIT), and one in central and anterior IT (cIT/aIT)^36^. Each image was presented approximately 50 times, using rapid visual stimulus presentation (RSVP). Each stimulus was presented for 100*ms*, followed by a mean gray stimulus interleaved between images. Each trial lasted 260*ms*. The image set consisted of 5120 images based on 64 object categories. These objects belonged to 8 high-level categories (tables, planes, fruits, faces, chairs, cars, boats, animals), each of which consisted of 8 unique objects. Each image consisted of a 2D projection of a 3D model added to a random background. The pose, size, and *x*- and *y*-position of the object was varied across the image set, whereby 2 levels of variation were used (corresponding to medium and high variation^36^). Multi-unit responses to these images were binned in 10*ms* windows, averaged across trials of the same image, and normalized to the average response to a blank image. This produced a set of 5120 images × 256 units × 25 timebins responses, which were the targets for our model features to predict. There were 88 units from V4, 88 units from pIT, and 80 units from cIT/aIT.

#### A.6.2 Fitting procedure

##### Generating train/test split

The 5120 images were split 75%-25% within each object category into a training set and a held-out testing set. All images were presented to the models for 10 time steps (corresponding to 100*ms*), followed by a mean gray stimulus for the remaining 15 time steps, to match the image presentation to the primates. The images are matched to the procedure when used to validate the models on ImageNet, namely they are bilinearly resized to 224 × 224 and normalized by the ImageNet mean ([0.485, 0.456, 0.406]) and standard deviation ([0.229, 0.224, 0.225]), applied per channel.

##### Model layer determination

We stipulated that units from each multi-unit array must be fit by features from a single model layer. To determine which one, we fit the features from the relevant feedforward BaseNet (either the 6-layer BaseNet or 11-layer BaseNet) to unit’s time-averaged response, and counted how many units had minimal loss for a given model layer, schematized in Step 2 of Figure 1. This yielded a mapping from the V4 array to model layer 3 of the 6-layer BaseNet and model layers 5 & 6 of the 11-layer BaseNet, pIT mapping to model layer 4 of the 6-layer BaseNet and model layers 7 & 8 of the 11-layer BaseNet, and cIT/aIT mapping to layer 5 of the 6-layer BaseNet and model layers 9 & 10 of the 11-layer BaseNet.

##### Mapping transform from models to neural responses

Model features from each image (i.e. the activations of units in a given model layer) were linearly fit to the neural responses by stochastic gradient descent with a standard L2 loss using a spatially factored mapping^55^, where each of the 256 units was fit independently. This spatially factored mapping is defined as follows: Given a model feature 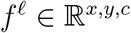 from layer ℓ, where *x* and *y* are the number of units in the spatial extent and *c* is the number of channels, we fit a spatial mask 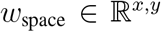 and a channel mask 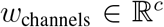 for each neuron *n* to predict the ground-truth neuron’s response *r_i,n,t_* at image *i* and timebin *t*. The predicted response can be written as:

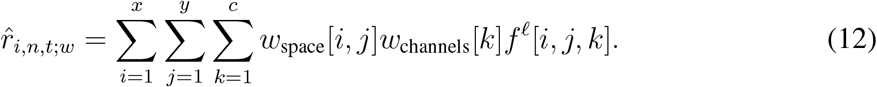

This mapping is implemented in the `factored_fc()` function of the `utils.cell_utils.py` file on our Github repository.

##### Loss function

After these layers were determined, model features were then fit to the entire set of 25 timebins for each unit using a shared linear model: that is, a single set of regression coefficients was used for all timebins, as schematized in Step 3 of Figure 1. The loss for this fitting was the average L2 loss across training images and 25 timebins for each unit, given by

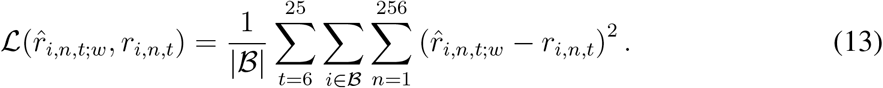

Note that *t* indexes model timesteps, which correspond to 10*ms* timebins, so *t* = 6 refers to the 60-70*ms* timebin, *t* = 7 refers to the 70-80*ms* timebin, and so forth.

We trained the temporally-fixed parameters *w* = [*w*_space_; *w*_channels_] of the mapping using the Adam optimizer^56^ with a learning rate of 1 × 10^−4^ and a training batch size 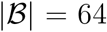 images. Additionally, we used a dropout^57^ level of 0.5 on the model features, prior to the mapping, as further regularization.

#### A.6.3 Metrics

To estimate a noise ceiling for each neuron’s response at each timebin, we computed the Spearman-Brown corrected split-half reliability *ρ_n_* of neuron *n*, averaged across 900 bootstrap iterations of split-half trials.

Let “Neural Predictivity” (used in Figure S5) refer to

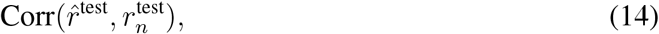

namely the Pearson correlation across test set images of the model’s response 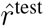 to the of any neuron *n*’s response 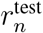 at a given timebin (or time-averaged).

The “Neural Predictivity (Noise Corrected)” (used in Figure 4 and Figure S7) for neuron n is given by

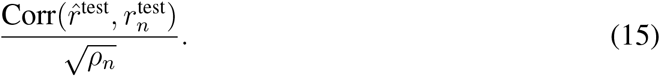

### A.7 Inter-animal consistency

We provide the definition and justification of the inter-animal consistency metric mentioned in Figure S7b. Suppose we have neural responses from two primates *A* and *B*. Let 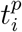 be the vector of true responses (either at a given timebin or averaged across a set of timebins) of primate *p* ∈ {*A, B*} on stimulus set *i* ∈ {train, test}. Of course, we only receive noisy observations of 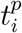, so let 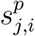 be the *j*-th set of *n* trials of 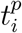. Finally, let *M*(*x*)_*i*_ be the predictions of a mapping *M* (e.g. PLS) when trained on input *x* and tested on stimulus set *i*. For example, 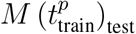 is the prediction of the mapping *M* on the test stimulus trained on the true neural responses from primate *p* on the train stimulus, and correspondingly, 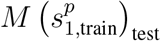 is the prediction of the mapping *M* on the test stimulus trained on the (trial-average) of noisy sample 1 on the train stimulus from primate *p*.

With these definitions in hand, the inter-animal mapping consistency from one primate *A* to another primate *B* corresponds to the following true quantity to be estimated:

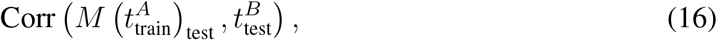

where Corr is the Pearson correlation across test stimuli. In what follows, we argue that this true quantity can be approximated with the following ratio of measurable quantities where we divide the noisy trial observations into two sets of equal samples:

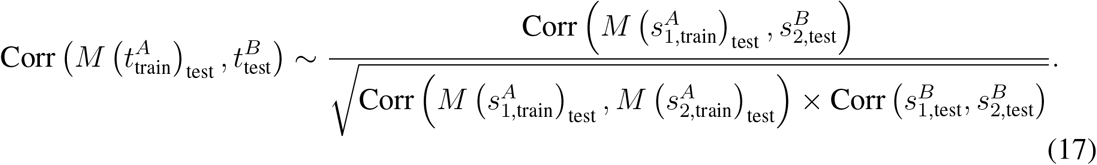

In words, the inter-animal consistency corresponds to the predictivity of the mapping on the test set stimuli from primate *A* to *B* on two different (averaged) halves of noisy trials, corrected by the square root of the mapping reliability on primate *A*’s test stimuli responses on two different halves of noisy trials and the internal consistency of primate *B*.

We justify the approximation in (17) by gradually eliminating the true quantities by their measurable estimates, starting from the original quantity in (16). First, we make the approximation that

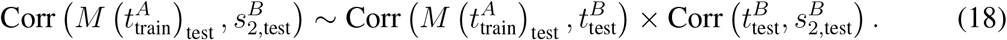

by transitivity of positive correlations (which is reasonable assumption when the number of stimuli is large). Next, by normality assumptions in the structure of the noisy estimates and since the number of trials (*n*) between the two sets is the same, we have that

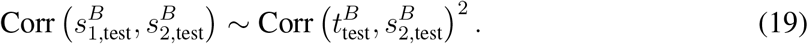

Namely, the correlation between the average of two sets of noisy observations of *n* trials each is approximately the square of the correlation between the true value and average of one set of *n* noisy trials. Therefore, from (18) and (19) it follows that

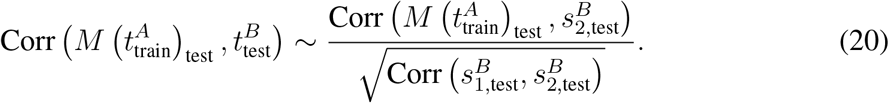

We have gotten rid of 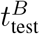, but we still need to get rid of the 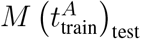 term. We apply the same two steps by analogy though these approximations may not always be true (though are true for additive Gaussian noise):

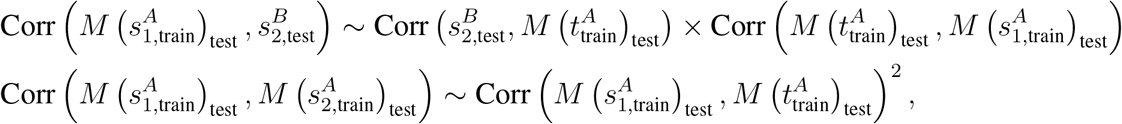

which taken together implies

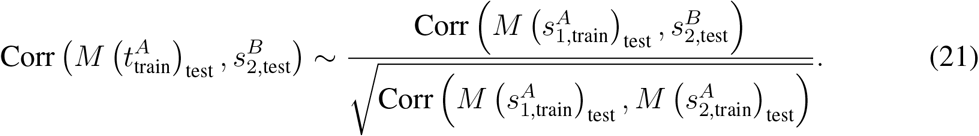

Equations (20) and (21) together imply the final estimated quantity given in (17).

### A.8 Object solution times (OSTs)

#### A.8.1 Generating model OSTs

Here we describe how we defined object solution times from both feedforward models and ConvRNNs. As depicted in Figure 3a, this is a multi-stage process that involves first identifying the most “IT-preferred” layers of each model.

##### Determining “IT-preferred” model layers

These are identified by a standard^34,12^ linear mapping using 25 component partial least squares regression (PLS), from model layer units to *time-averaged* IT (namely, pIT/cIT/aIT) responses from the neural data described in Section A.6.1, and corroborates the results obtained by the same procedure described in Section A.6.2. We use this neural data as it has both V4 and IT responses, and demonstrates a disjoint set of layers between the preferred V4 model layers and preferred IT layers.

##### Mapping model timepoints to IT timepoints

Once these “IT-preferred” model layers are identified, we then map these model timepoints to 10*ms* timebins as in the IT data. For ConvRNNs with intrinsic temporal dynamics, this mapping is one-to-one, we simply concatenate the model layers at each timepoint to construct an entire IT pseudopopulation, and each timepoint of the ConvRNN corresponds to a 10*ms* timebin between 70-260*ms*. For feedforward models, we map each “IT-preferred” layer to a 10*ms* timebin between 70-260*ms*. If the number of “IT-preferred” layers for a feedforward model matches the total number of timebins (19), then there is only one admissible mapping, corresponding to the “uniform” mapping, whereby the earliest (in the feedforward hierarchy) layer is matched to the earliest 10*ms* timebin of 70*ms*, and so forth. On the other hand, if the number of “IT-preferred” layers is strictly less than the total number of timebins, then we additionally consider a “graded” mapping that picks a random sample of units from one layer to the next so that the number of feedforward layers exactly matches the total number of timebins.

##### Obtaining model d′ values

Once a timepoint mapping is selected, we compute the model object solution time (OST) in the same manner as the OST is computed for IT12. Specifically, we train an SVM (*C* = 5 × 10^4^) separately for each model timepoint after it has been dimension reduced through PCA (with 1000 components) to solve the ten-way categorization task for each image. The ten categories are apple, bear, bird, car, chair, dog, elephant, person, plane, and zebra. 1000 images constitute the training set of the SVM (100 images per category) and 320 images are randomly chosen to be in the test set. We perform 20 trials each of 10 train/test splits to get errorbars, where each image is in the test set at least once. The model *d*′ for that image is computed in the same manner as previously done for the ground truth IT response *d*′ (see Kar et al. 2019^12^ for details), only being computed from the SVM when it has been in the test set and is bounded between −5 and 5. Since this dataset consists of 1320 grayscale images presented centrally to behaving primates for 100*ms*, there are therefore 1320 *d*′ values (one for each image) for any given model, constituting its “I1” vector^31^.

##### Correlating model OST with IT OST

The OST of the model therefore is the first model timepoint in which the *d*′ reaches the recorded primate *d*′ for that image, as was previously done to compute the ground truth IT OST12. Using the Levenberg–Marquardt algorithm, we further linearly interpolate between 10*ms* bins to determine the precise millisecond that the response surpassed the primate’s behavioral output for that image (as was done analogously with the IT population’s OST). Finally, we compare the model OST to the IT OST via a Spearman correlation across the common set of images solved by *both* the model and IT.

#### A.8.2 Relating the linear mapping to neural responses with the OST behavioral metric

The IT population OST was computed from primarily anterior IT (aIT) responses^12^. Therefore, to isolate the interaction a linear mapping of model features to neural responses (as we do in neural response prediction described in Section A.6) might have compared to directly computing the OST from the original model features, we turned to neural data collected from 486 aIT units on 1100 greyscale images.

For each model, we train a linear mapping on this dataset, with 550 images used for training the mapping and 550 images are held-out for the test set. We observe similar conclusions as with the original neural data in Section A.6 for both the temporally-fixed linear mapping in Figure S6 (in the “aIT” panel), and with a temporally-varying PLS mapping in Figure S7 (“aIT” in panel (a) as well as the data used in panel (b)), all from layer 10 of the 11-layer BaseNet/ConvRNNs.

With these observations, we then proceeded to evaluate the effect of the linear mapping on OST correlations in Figure S3. Crucially, in this setting, we train a 100 component PLS mapping on the 526 images for which an IT *d*′ is *not* defined, in order to ensure that the images from Section A.8.1 that the OST correlation is evaluated on are not the same images the PLS mapping was trained with.

## Extended Data

## Supplementary Figures

**Figure S1:**
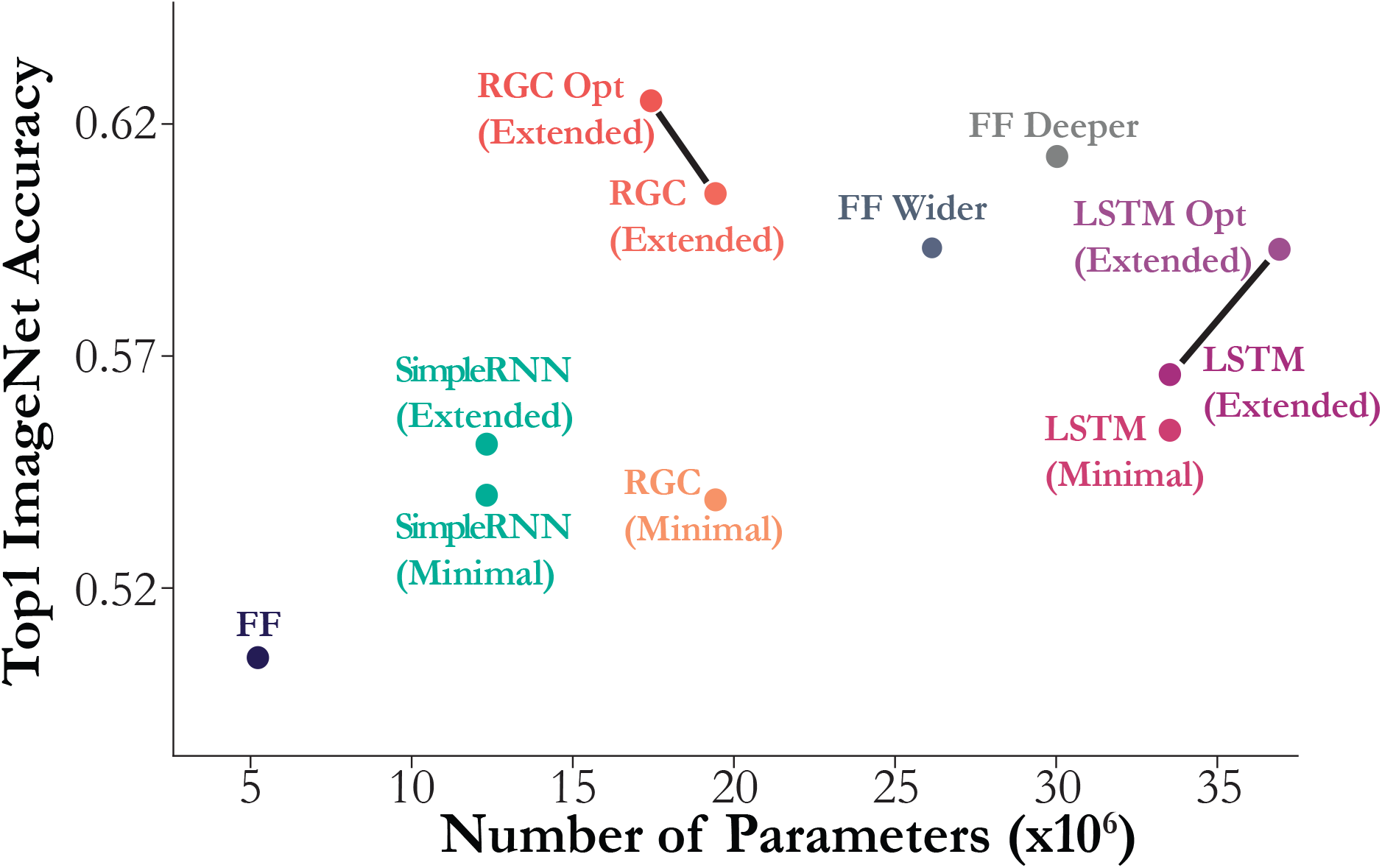
Performance of shallow ConvRNN and feedforward models as a function of number of parameters. Colored points incorporate the respective ConvRNN circuit into the shallow, 6-layer feedforward BaseNet architecture (“FF”). “Minimal” is defined as the minimum number of timesteps (7) after the initial feedforward pass whereby all recurrence connections were engaged at least once, which the model was trained with. “Extended” is a greater number of timesteps (16) that the model was trained for given optimization and memory constraints. Hyperparameter-optimized versions of the LSTM (“LSTM Opt”) and Reciprocal Gated Circuit ConvRNNs (“RGC Opt”) are connected to their non-optimized versions by black lines. Note that the feedforward (FF) models are already optimized for the relevant hyperparameters of batch size, learning rate, and L2 regularization. The SimpleRNN is also hyperparameter optimized since unlike the more sophisticated ConvRNN circuit architectures of the LSTM and RGC, it is unable to train otherwise – with layer normalization being an important factor (see Section A.3.2 for more details).

**Figure S2:**
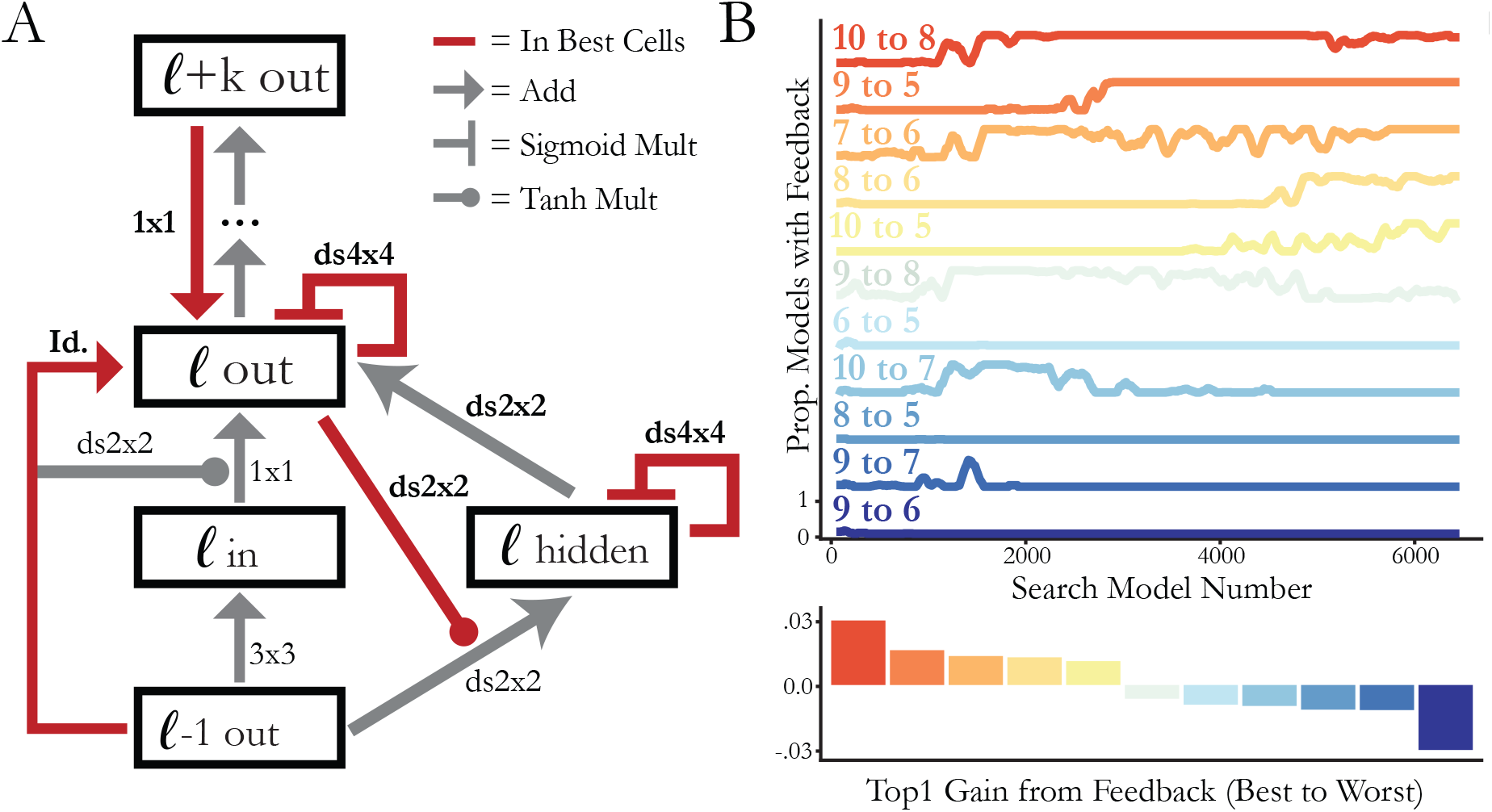
Optimal local recurrent circuit motif and global feedback connectivity. **(a) RNN circuit structure from the top-performing search model.** Red lines indicate that this hyperparameter choice (connection and filter size) was chosen in each of the top unique models from the search. *K* × *K* denotes a convolution and ds *K* × *K* denotes a depth-separable convolution with filter size *K* × *K*. **(b) Long-range feedback connections from the search.** (Top) Each trace shows the proportion of models in a 100-sample window that have a particular feedback connection. (Bottom) Each bar indicates the difference between the median performance of models with a given feedback and the median performance of models without that feedback. Colors correspond to the same feedback connectivity as above.

**Figure S3:**
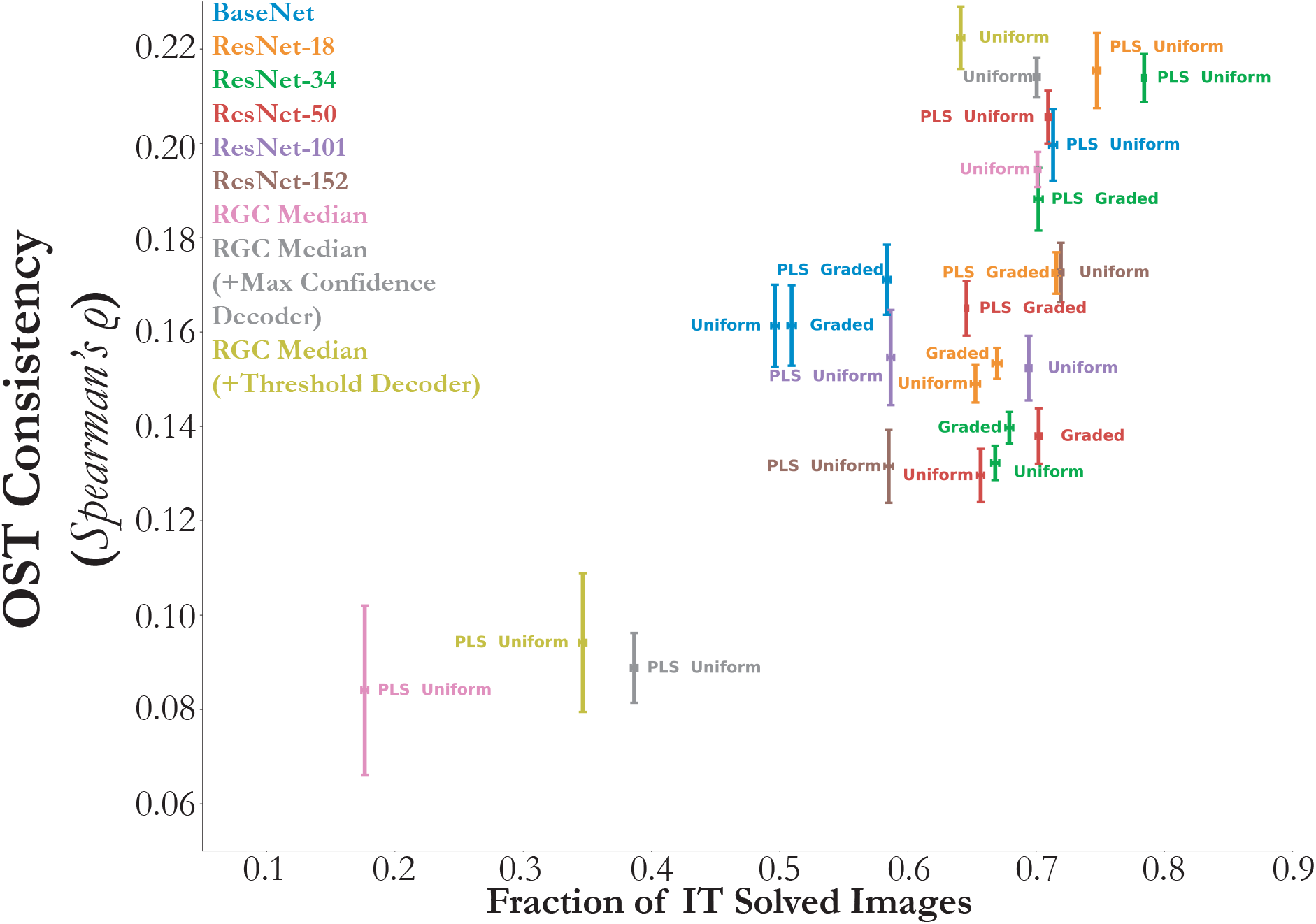
Behaviorally harmful effect of dimensionality reduction due to linear transform. Mean and s.e.m. are computed across train/test splits (*N* = 10) when that image (of 1320 images) was a test-set image, with the Spearman correlation computed with the IT solution times across the imageset mutually solved by the given model and IT. As can be seen, a temporally-graded mapping directly from the model features of feedforward models always attains OST consistency at least that of the uniform one (“Graded” vs. “Uniform” comparison). We additionally train a 100 component PLS regression to IT responses at each defined model timepoint, where the responses are to a *different* set of images than used to evaluate the OST metric. This procedure, detailed in Section A.8.2, results in an image-computable model on which the OST metric is evaluated on and corresponds to “PLS” prepended to the name of each point on this plot, for any given model and associated temporal mapping. As can be seen, “PLS Uniform” for the BaseNet and ResNet-34 match the OST consistency of the RGC Median ConvRNNs from their original model features. However, “PLS Uniform” for the ConvRNNs and ResNet-101 and ResNet-152 have a significant decrease in OST consistency compared to when evaluated on their original model features, indicating the behaviorally harmful effect of dimensionality reduction due to PLS.

**Figure S4:**
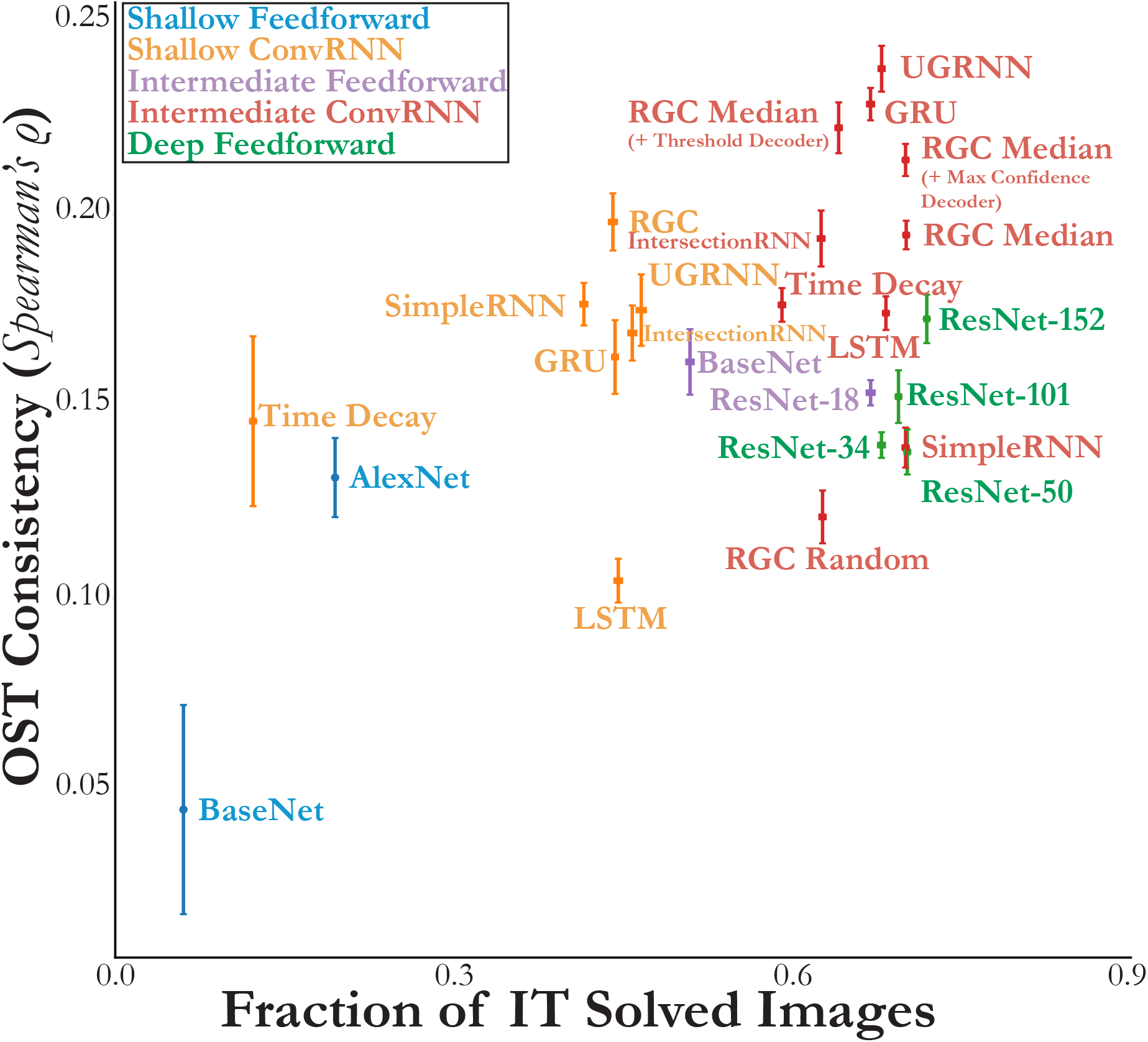
Decoding strategy can impact IT OST. Same as Figure 3d, but we additionally embed decoders to the Reciprocal Gated Circuits (RGC), see definitions in Section A.5. Mean and s.e.m. are computed across train/test splits (*N* = 10) when that image (of 1320 images) was a test-set image, with the Spearman correlation computed with the IT object solution times (analogously computed from the IT population responses) across the imageset solved by both the given model and IT, constituting the “Fraction of IT Solved Images” on the *x*-axis. We start with either a shallow base feedforward model consisting of 5 convolutional layers and 1 layer of readout (“BaseNet” in blue) as well as an intermediate-depth variant with 10 feedforward layers and 1 layer of readout (“BaseNet” in green), detailed in Section A.2.1. From these base feedforward models, we embed recurrent circuits, resulting in either “Shallow ConvRNNs” or “Intermediate ConvRNNs”, respectively.

**Figure S5:**
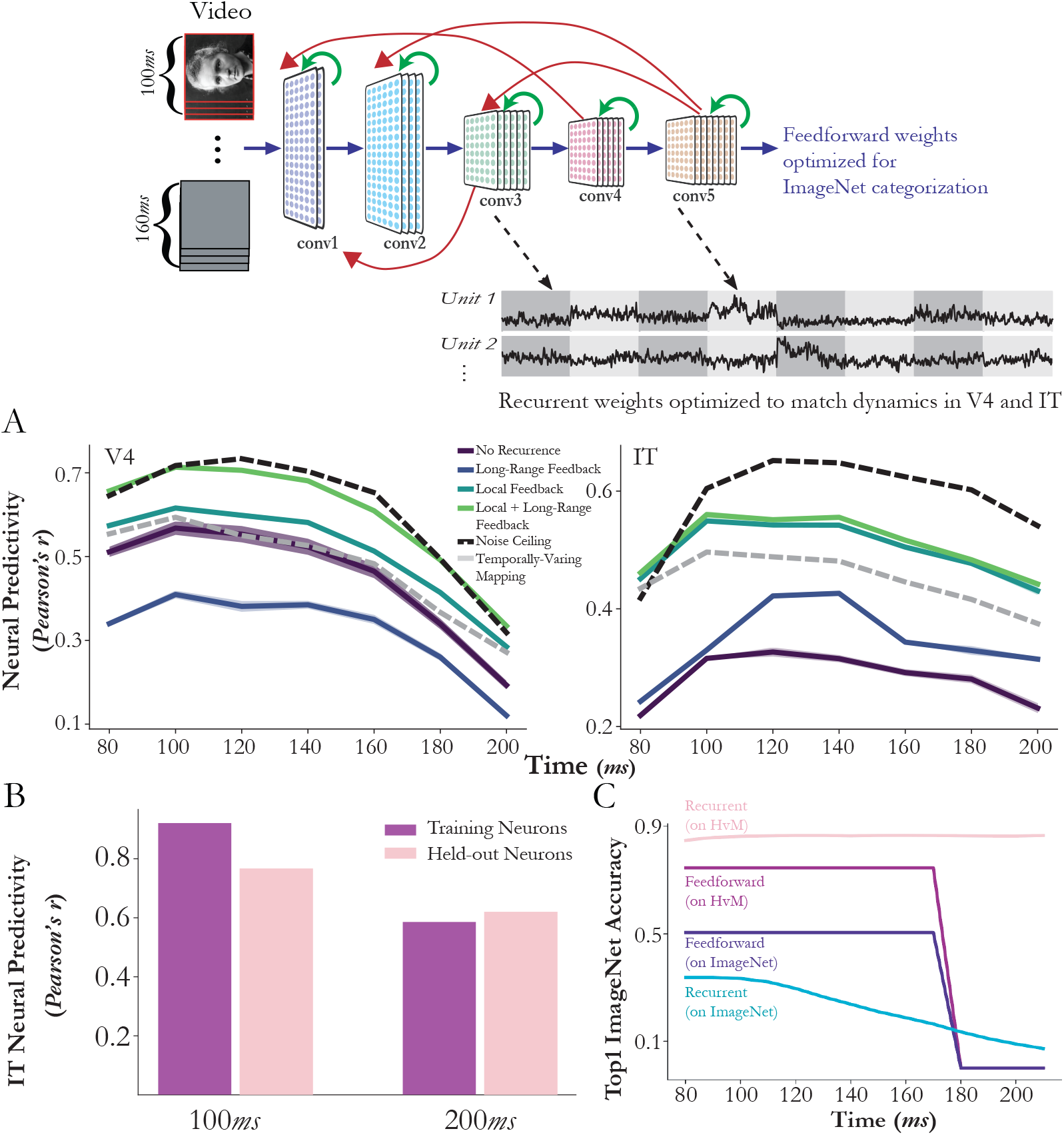
**(a) Both local recurrence and global feedback are needed to best fit neural data.** Among a wide range of architectures with different local recurrent motifs and global feedback patterns, the best architecture was one with both gated local recurrence and a global feedback. Local recurrent circuits were particularly useful for improving fits to IT neurons (*N* = 168), whereas both local recurrence and global feedback were critical for improving fits to V4 neurons (*N* = 88). Except for “temporally-varying mapping”, fixed model-unit-to-neuron linear mappings were fixed across all time bins, constraining trajectories to be produced by actual dynamics of the network. In contrast, “temporally-varying mapping” indicates an independent PLS regression for each time bin. The fact that models with local recurrence and global feedback are better than “temporally-varying mapping” suggests that some nonlinear dynamics at earlier layers contributed meaningfully to network fits. S.e.m. across four splits of held-out test images. **(b) Held-out neural predictivity.** At both 100*ms* and 200*ms*, this direct fitting procedure to the dynamics generalizes to neurons held-out (right bars) in the fitting procedure, a stronger test of generalization than held-out images depicted in the left bars. **(c) Underfitting to the task.** However, a subtle overfitting to the neural image distribution occurs, whereby the task-optimized network whose dynamics are trained on the V4 and IT neural dynamics no longer transfers to ImageNet.

**Figure S6:**
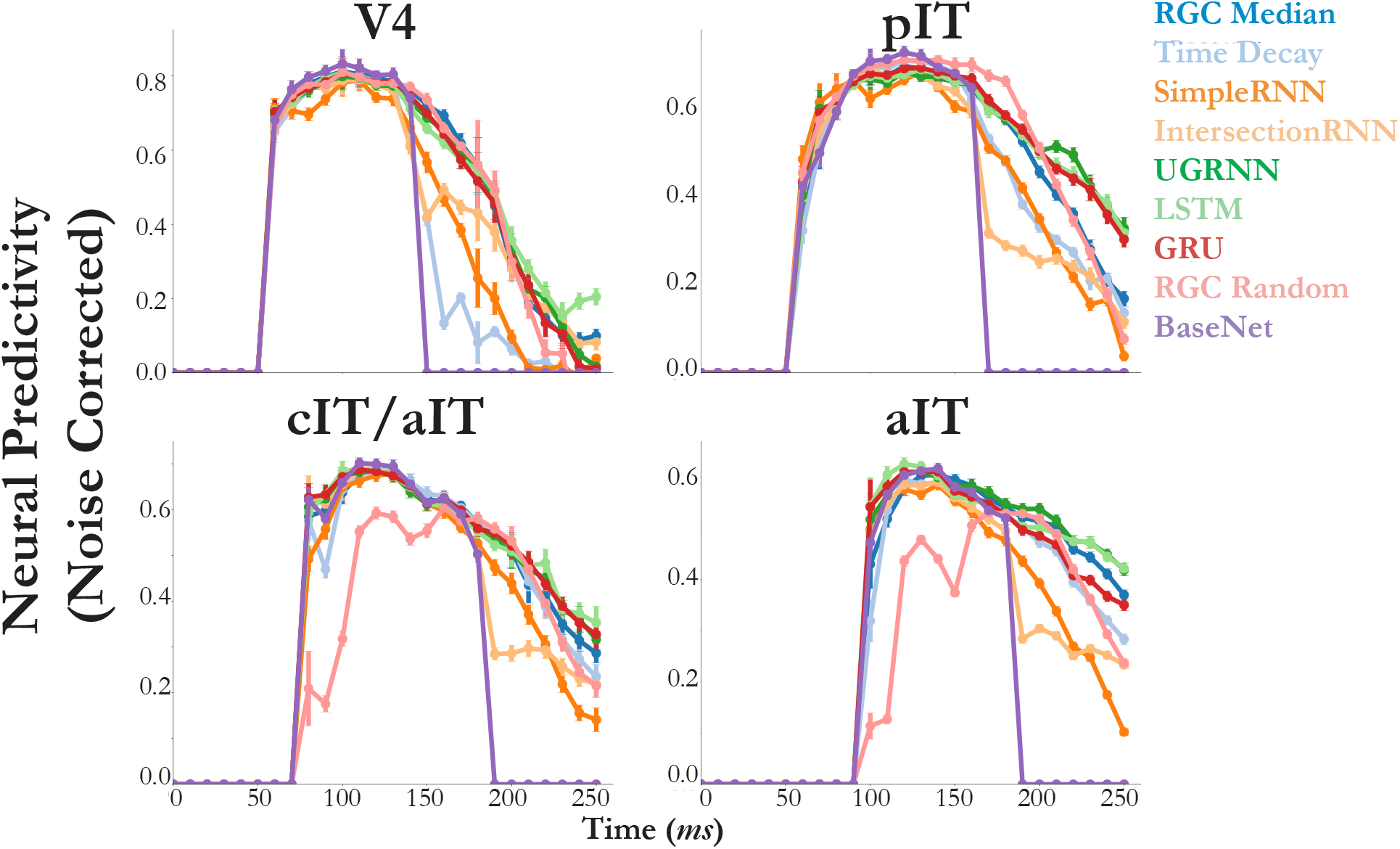
Intermediate ConvRNN circuits are differentiated by primate ventral stream neural dynamics. Fitting model features of ConvRNNs with a temporally-fixed linear mapping to neural dynamics approaches the noise ceiling of these responses in most cases. The y-axis indicates the median across neurons of the explained variance between predictions and ground-truth responses on held-out images. Error bars indicates the s.e.m across neurons (*N* = 88 for V4, *N* = 88 for pIT, *N* = 80 for cIT/aIT, and *N* = 486 for aIT). Note that “aIT” refers to a separate neural dataset from primarily anterior IT neurons, detailed in Section A.8.2. The onset time of the response is the first timepoint the area-preferred model layer (see Section A.6.2 for details) of the base feedforward model (“BaseNet”), which all these circuits share, receives its input. As can be seen, the feedforward BaseNet model (purple) is incapable of generating a response beyond the feedforward pass, and certain types of ConvRNN circuits added to the feedforward model are less predictive than others.

**Figure S7:**
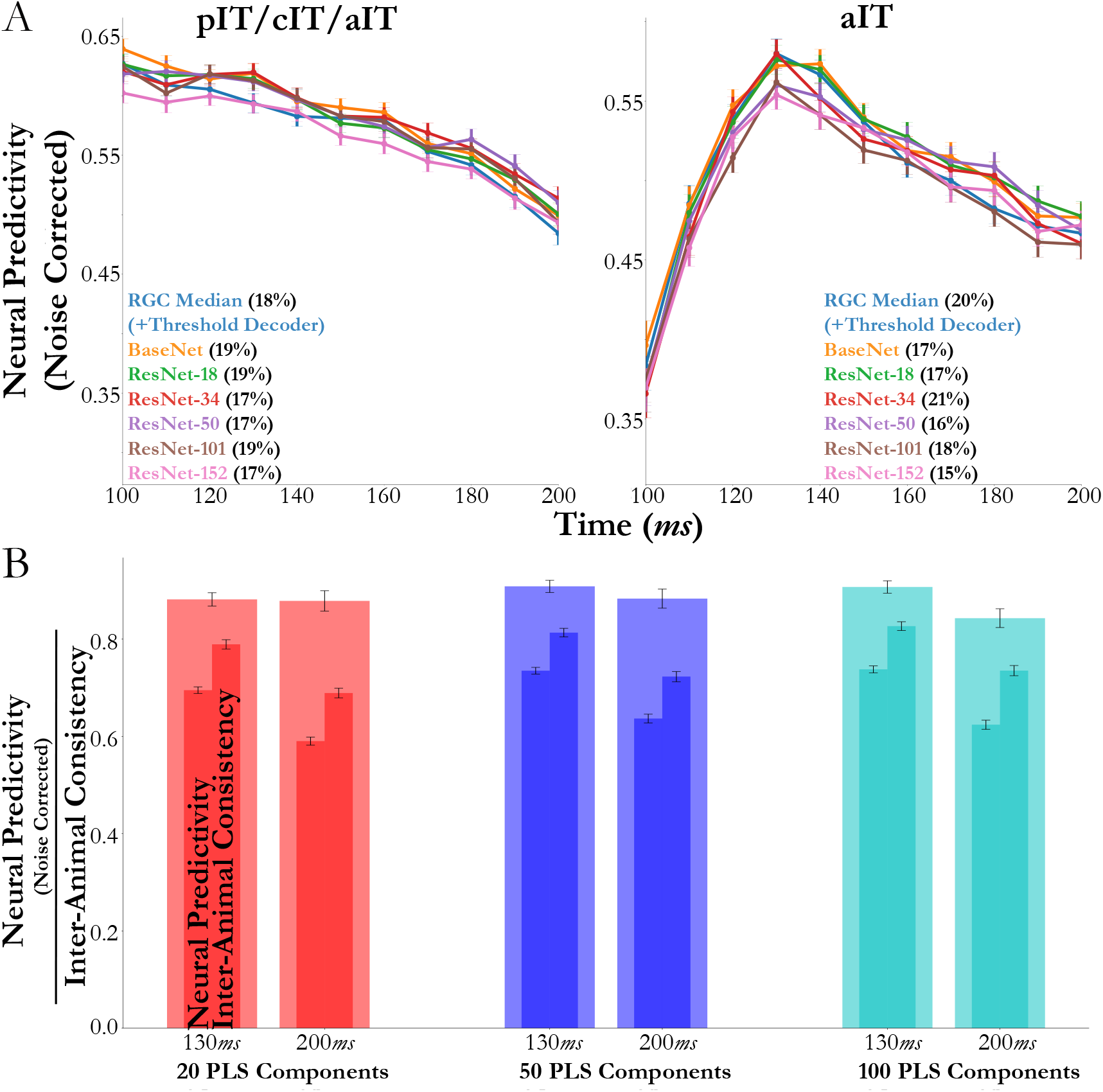
**(a) Increasing feedforward depth does *not* account for drop in median explained variance from early to late timepoints.** We observe a similar drop in median explained variance from 130-140*ms* to 200-210*ms*, between the intermediate-depth ConvRNN and deeper feedforward models, where we fix each model’s training image size and batch size to be able compare across depths. To compare these two models, we subselect for high reliability neurons (above 0.3 split-half consistency) and use a temporally-varying mapping (PLS 25 components). Note that the temporally-varying mapping implies providing the feedforward models with a constant input stream (unlike the primates and ConvRNNs, which are given a 100*ms* presentation) in order for them to produce a (constant) response at every timepoints. We plot the median and s.e.m. predictivity in both panels per timebin (*N* = 108, 113, 117, 123, 118, 118, 116, 115, 108, 99, 86 neurons for each timebin in the “pIT/cIT/aIT” panel, and *N* = 247, 313, 378, 441, 437, 411, 397, 391, 392, 384, 380 neurons for each timebin in the “aIT” panel). **(b) Drop in explained variance may be exhibited in inter-animal consistency.** Using the neural data described in Section A.8.2, we see a similar inter-animal consistency (metric detailed in Section A.7) at 130-140*ms* and 200-210*ms*, as we do with the 11-layer BaseNet. Median and s.e.m. across aIT neurons (*N* = 441 at 130-140*ms* and *N* = 380 at 200-210*ms*) from the dataset described in Section A.8.2.

## Footnotes

a Mean OST difference 0.0120 and s.e.m. 0.0045, Wilcoxon test on uniform vs. graded mapping OST consistencies across feedforward models, *p* < 0.001; see also Figure S3.

b Paired *t*-test with Bonferroni correction: shallow Time Decay vs. “BaseNet” in blue, mean OST difference 0.101 and s.e.m. 0.0313, *t*(9) ≈ 3.23, *p* < 0.025; intermediate Time Decay vs. “BaseNet” in purple, mean OST difference 0.0148 and s.e.m. 0.00857, *t*(9) ≈ 1.73, *p* ≈ 0.11.

c Paired *t*-test with Bonferroni correction: shallow RGC vs. “BaseNet” in blue, mean OST difference 0.153 and s.e.m. 0.0252, *t*(9) ≈ 6.08, *p* < 0.001; intermediate UGRNN vs. ResNet-152, mean OST difference 0.0652 and s.e.m. 0.00863, *t*(9) ≈ 7.55, *p* < 0.001; intermediate GRU vs. ResNet-152, mean OST difference 0.0559 and s.e.m. 0.00725, *t*(9) ≈ 7.71, *p* < 0.001; RGC Median vs. ResNet-152, mean OST difference 0.0218 and s.e.m. 0.00637, *t*(9) ≈ 3.44, *p* < 0.01.

d Paired *t*-test with Bonferroni correction, mean OST difference 0.0195 and s.e.m. 0.00432, *t*(9) ≈ –4.52, *p* < 0.01.

e Paired *t*-test with Bonferroni correction, mean OST difference 0.0279 and s.e.m. 0.00634, *t*(9) ≈ –4.41, *p* < 0.01.

f Wilcoxon test with Bonferroni correction *p* < 0.001 for each ConvRNN vs. Time Decay, except for the SimpleRNN *p* ≈ 0.46 for pIT.

g Wilcoxon test with Bonferroni correction between each of these ConvRNNs vs. the SimpleRNN on late phase dynamics, *p* < 0.001 per visual area.

h Paired *t*-test with Bonferroni correction: RGC Median vs. PLS Uniform BaseNet, mean OST difference −0.0052 and s.e.m. 0.0061, *t*(9) ≈ –0.86, *p* ≈ 0.41; RGC Median with Threshold Decoder vs. PLS Uniform ResNet-18, mean OST difference 0.00697 and s.e.m. 0.0085, *t*(9) ≈ 0.82, *p* ≈ 0.43; RGC Median with Max Confidence Decoder vs. PLS Uniform ResNet-34, mean OST difference 0.0001 and s.e.m. 0.0079, *t*(9) ≈ 0.02, *p* ≈ 0.99.

## Notes

https://github.com/neuroailab/convrnns

### Competing Interest Statement

The authors have declared no competing interest.

### Summary of Updates

Final version accepted for publication in Neural Computation

